# Transient Poly(ADP-Ribose) Triggers FUS Condensation Hysteresis via a Prion-Like Mechanism

**DOI:** 10.1101/2025.07.03.659157

**Authors:** Hongrui Liu, Yuxuan Cai, Leilei Shi, Meenakshi Pillai, Nilimesh Das, Haley E. Tarbox, Yingda Ge, Kun Yue, Xingyi Yang, Piyush Rath, Mohsen Badiee, Charina S. Fabilane, Jamie B. Spangler, Mark T. Bedford, Sua Myong, Stephen D. Fried, Xinqiang Ding, Anthony K. L. Leung

## Abstract

Hysteresis—where a system retains memory of a transient stimulus—is common in signaling but can also arise in intracellular organization. DNA repair foci, a type of biomolecular condensate, are initiated by the short-lived noncanonical nucleic acid poly(ADP-ribose) (PAR). PAR recruits proteins with prion-like domains (PrLDs), such as Fused in Sarcoma (FUS), and initiates their condensation, which persists even after PAR degradation. How FUS transitions from PAR-dependent to PAR-independent condensation remains unclear. Here, we show that PAR binding triggers a conformational switch in FUS, enabling sustained condensation. PAR binds to the C-terminal arginine-rich region of FUS, displacing intramolecular contacts, and exposing the N-terminal PrLD. This conformational opening allows PrLD interactions *in trans*, stabilizing condensates independently of PAR. FUS thus undergoes a regulated, nucleated conformational conversion—reminiscent of classical prions. This mechanism implies a paradigm of nucleic acid-induced conformational memory that may underlie hysteresis in intracellular organization in health and disease.

**HIGHLIGHTS:**

● PAR-initiated FUS condensation follows a bi-modular mechanism involving both FUS termini.
● Simulations predict and experiments confirm PAR disrupts FUS intramolecular contacts.
● Upon condensation, FUS adopts a conformation with its N-terminus open for interactions.
● N-terminal interactions maintain FUS condensation when PAR degrades during DNA repair.

## INTRODUCTION

Many biological systems exhibit hysteresis, wherein a transient stimulus induces an altered state that persists after the initiating signal has dissipated^1,2^. This mechanism enables cells to maintain functionally distinct states in response to short-lived inputs. Hysteresis is well-documented in signaling contexts, such as cell fate decision, immune responses and neural circuits, and it is typically driven by feedback loops^3^. Beyond molecular signaling, emerging evidence suggests that hysteresis can also arise at the mesoscopic scale. For instance, *in vitro* experiments have shown that nucleic acids can trigger proteins involved in condensate or filament formation to assemble into self-sustaining structures that persist even after the nucleic acid scaffold is removed^4–7^. However, the underlying molecular mechanisms and the extent to which this phenomenon occurs in native biological contexts remain unclear.

Nucleic acids, including DNA^8–11^, RNA^12–14^, and the noncanonical nucleic acid poly(ADP-ribose) (PAR)^15,16^, often serve as multivalent structural scaffolds for mesoscopic assemblies across diverse biological processes^17–19^. For example, mRNA and PAR are critical for assembling stress-induced condensates called stress granules; their continual presence sustains these structures, while their degradation leads to condensate dissolution^20,21^. Yet nucleic acids can also act transiently, serving as short-lived triggers rather than structural scaffolds^4–6^. A compelling example in cells is DNA repair foci, a type of inducible biomolecular condensate that compartmentalizes and orchestrates the DNA repair process^9,22,23^. Within seconds of DNA damage, the PAR-synthesizing enzyme PARP1^24^ localizes to DNA breaks and synthesizes PAR^25–27^, which serves as a transient scaffold that recruits effector proteins^27–29^—including the FET family members FUS (Fused in Sarcoma) and EWSR1 (Ewing’s Sarcoma RNA-Binding Protein 1)—to drive repair foci formation^22,23,30–32^ **(Figure 1A)**. Notably, FDA-approved PARP1 inhibitors, which block PAR synthesis, prevent repair foci formation, highlighting the necessity of PAR in their initiation and its therapeutic significance^22,23,29,33,34^. However, within 1–2 min of its synthesis, PAR is rapidly degraded by poly(ADP-ribose) glycohydrolase (PARG), transitioning the repair process to a PAR-independent phase^35^ **(Figure 1A)**. Yet, repair foci persist for more than 10 min^36^. This sustained presence exemplifies hysteresis, in contrast to other nucleic acid-seeded condensates which rapidly disassemble upon degradation of nucleic acid scaffolds^8,20^. If PAR is essential for initiating repair foci but is rapidly degraded, what sustains these condensates over extended periods?

**Figure 1.**
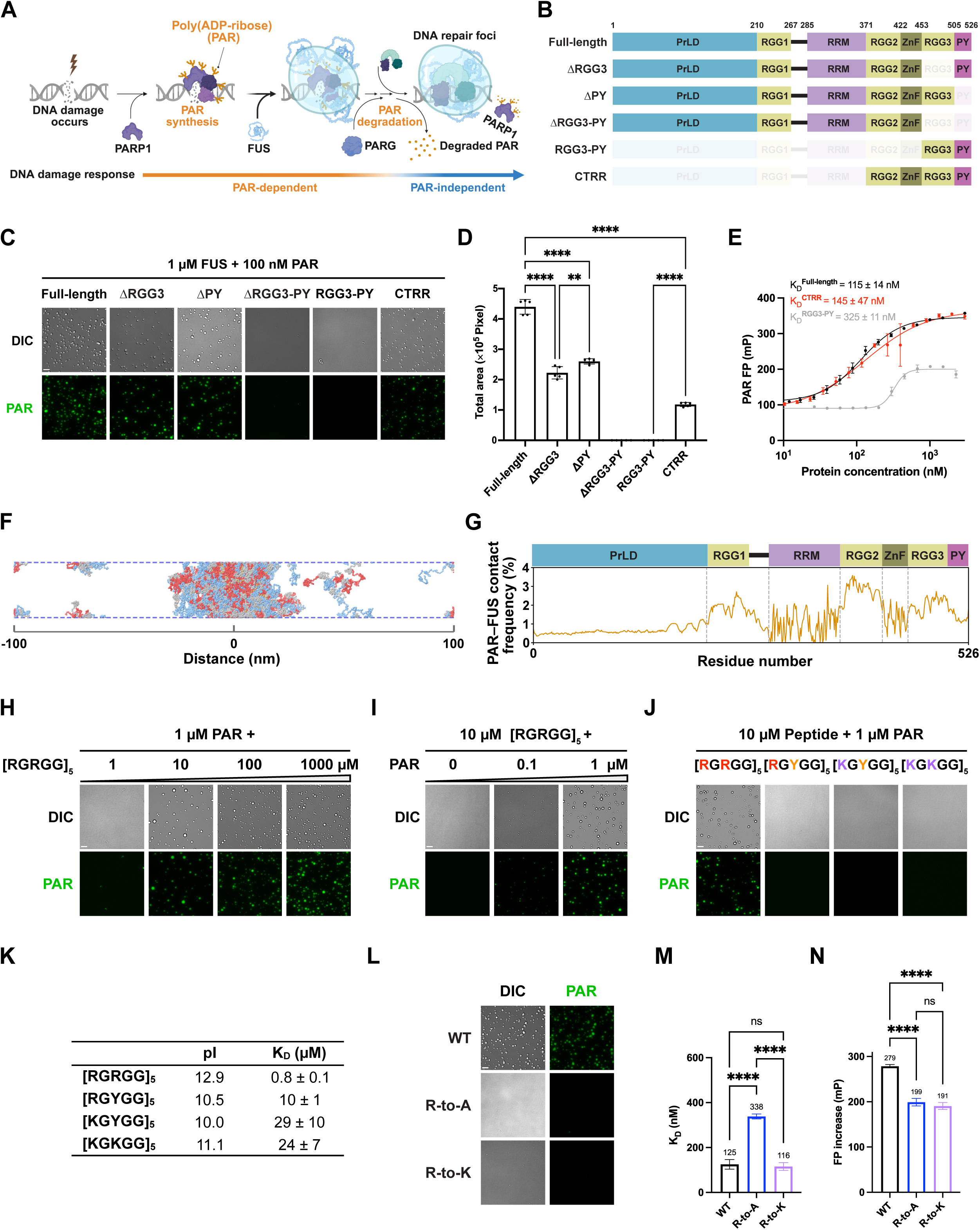
The CTRR of FUS Enables PAR-Driven Condensation Through Arginine-Mediated Interactions Beyond Simple Charge Complementarity. **(A)** Schematic of PAR-initiated FUS condensation during DNA repair foci formation. The process transitions from a PAR-dependent phase, in which PAR synthesized by PARP1 triggers FUS condensation, to a PAR-independent phase, where FUS condensates persist even after PARG degrades PAR. **(B)** Domain structures of full-length FUS and its truncation mutants. **(C)** Representative microscopic images of condensation results for 1 µM full-length FUS or its truncation variants with 100 nM 16-mer PAR under physiological salt concentrations from DIC and fluorescence microscopy showing Cy3-PAR signals. n = 5. Scale bar, 5 µm. **(D)** Quantification of the total area covered by condensates formed in (C). n = 5; error bars, SD; ordinary one-way ANOVA with Tukey’s multiple comparisons test, p < 0.01 (**) and p < 0.0001 (****). **(E)** Fluorescence anisotropy assays of full-length FUS (black) and the RGG3-PY (gray) and CTRR (red) fragments with 3 nM AF488-labeled PAR. Fluorescence polarization (FP) values of AF488-PAR are plotted and fitted using the Hill equation. n = 3; error bars, SD. See also Figure S1B. **(F)** A configurational snapshot of FUS slab simulations with 16-mer PAR at a simulation temperature of 380 K, which enables efficient sampling of conformations within tractable computational timescales (See **Methods**). PrLD is shown in blue, RGG1-RRM in gray, and the remaining C-terminal region (later referred to as CTRR) is in red; PAR is depicted in amber. **(G)** Relative contact frequency between PAR and FUS in slab simulations as in (F) (See **Methods**). **(H)** Representative microscopic images showing condensation results for 1 µM 16-mer PAR with a titration of [RGRGG]_5_ peptide. n = 5. Scale bar, 5 µm. See also Figure S1F. **(I)** Representative microscopic images showing condensation results for 10 µM [RGRGG]_5_ peptide with either 0.1 or 1 µM PAR. n = 5. Scale bar, 5 µm. See also Figure S1G. **(J)** Representative microscopic images showing condensation results for 10 µM [RGRGG]_5_ peptide or its arginine-substituted derivatives with 1 µM PAR. n = 5. Scale bar, 5 µm. See also Figure S1I. **(K)** Table showing the PAR-binding affinity of the [RGRGG]_5_ peptide and its arginine-substituted derivatives, measured by electrophoretic mobility shift assay (EMSA). Isoelectric points (pI) are indicated for each peptide. Equilibrium dissociation constants (K_D_) were determined by fitting EMSA data to the Hill equation. Values represent mean ± SD, n = 3. See also Figures S1J and S1K. **(L)** Representative microscopic images of condensation results for 1 µM wild-type FUS or its arginine mutants (R-to-A and R-to-K) with 100 nM PAR. n = 5. **(M, N)** Comparisons of K_D_ values and the increase in FP from fluorescence anisotropy assays of wild-type FUS and its arginine mutants with 3 nM AF488-labeled PAR. n = 3; error bars, SD; ordinary one-way ANOVA with Tukey’s multiple comparisons test, with significance levels: not significant (ns), p < 0.05 (*), and p < 0.0001 (****). See also Figure S1N.

Recent evidence indicates that FUS serves as a key component in DNA repair foci formation. FUS is a multifunctional, multi-domain nucleic acid-binding protein with extensive intrinsically disordered regions^31,37,38^. As one of the earliest proteins to respond to DNA damage^37,39^, FUS is recruited to DNA damage sites and undergoes condensation in a PAR-dependent manner^22,31–33,37^, helping maintain the PARP1–DNA co-condensation and organize repair machinery for efficient DNA repair^9,32,37–40^.

Notably, FUS condensation persists at DNA damage sites for over 5–10 min^22,32,37–39^, even after PAR degradation^41^. *In vitro*, FUS condenses on its own at high concentrations (≥ 3–5 µM)^33,42,43^, but as little as 1 nM PAR can trigger FUS condensation at physiologically relevant levels (∼1 µM)^4,33^. Importantly, these PAR-initiated FUS condensates persist even after PAR degradation by PARG^4^. These data suggest that FUS facilitates a transition from a PAR-dependent to a PAR-independent state in DNA repair foci **(Figure 1A)**. Yet, the molecular mechanisms orchestrating this hysteretic transition remain unclear: How does PAR initially trigger FUS condensation, and how is condensate integrity sustained once the initiating scaffold is lost?

Here, we reveal a bi-modular mechanism in which the intrinsically disordered regions at both termini of FUS coordinate to mediate the transition from a PAR-driven to a protein-driven condensate. PAR interacts with the arginine-glycine-rich C-terminal region of FUS, triggering condensation and disrupting *cis* interactions between the C- and N-terminal regions of FUS. This conformational change exposes the N-terminal prion-like domain (PrLD) of FUS, enabling interactions *in trans* that sustain FUS condensation even after PAR degradation. This coordinated bi-modular mechanism bears a striking resemblance to nucleated conformational conversion^44^—a model originally proposed to explain prion propagation, in which a transient seed imprints a self-perpetuating conformation. In this context, PAR functions not only as a trigger but also as a transient scaffold that nucleates a structurally permissive intermediate, allowing the PrLD of FUS to propagate a persistent condensate state. In this mechanism, PrLD acts as a molecular memory element, stabilizing the condensate even after the initiating nucleic acid signal is lost. This work thus establishes a new paradigm for condensation hysteresis, in which transient nucleic acid signals initiate a spatial state that is subsequently maintained through PrLD interactions.

## RESULTS

### The C-Terminal RGG-Rich Region (CTRR) of FUS Is Sufficient to Interact and Condense with PAR

To elucidate the molecular basis for FUS condensation induced by transient PAR signaling, we first examined the protein domains required for this process. In cells, the PAR-dependent enrichment of FUS at DNA damage sites is mediated by the RGG3-PY region at its C-terminus, which contains an arginine-glycine-glycine (RGG) repeat followed by a proline-tyrosine (PY)-type nuclear localization sequence^22,32^. To determine whether RGG3-PY is critical for PAR-initiated condensation, we generated truncation mutants and performed condensation assays *in vitro* **(Figure 1B)**. Briefly, purified recombinant full-length and truncated FUS (1 µM) were combined with fluorescently labeled PAR (100 nM) under physiological salt concentrations *in vitro* **(Figure 1C)**.

Deleting either the RGG3 or PY regions significantly reduced FUS–PAR condensation, as quantified by the total condensate area within fixed-sized fields across multiple replicates **(Figures 1B–1D and S1A**). Deleting both regions abolished condensation entirely under the same conditions. We infer that the RGG3-PY region is necessary for FUS–PAR condensation. Notably, the RGG3-PY fragment alone exhibited a threefold weaker PAR-binding affinity compared to full-length FUS, based on fluorescence anisotropy studies **(Figures 1B, 1E, S1A and S1B),** and this fragment was insufficient to form condensates with PAR **(Figures 1C and 1D)**. Therefore, additional regions of FUS work alongside RGG3-PY to promote PAR-initiated condensation.

To identify additional FUS regions that interact with PAR during condensation, we utilized molecular dynamics simulations to capture interactions critical for phase separation. FUS was modeled using two well-established coarse-grained approaches: a native-structure-based Gō-like model for its structured domains and an updated hydrophobicity scale (HPS) model for its disordered regions (see **Methods**)^45–50^, with each residue represented by one particle in both models. Additionally, we developed a custom HPS model for PAR, based on end-to-end distances measured in single-molecule Förster resonance energy transfer (smFRET) experiments^51^. Key parameters were optimized using FRET data for PAR of various lengths (see **Methods**), with simulation outputs closely matching experimental results **(Figure S1C)**^52^.

We performed slab simulations with 100 FUS and 10 PAR molecules in an elongated periodic box **(Figure 1F)**, designed to emulate FUS–PAR condensation while maintaining the experimental molar ratio. For comparison, FUS-only condensation was also simulated. Both systems exhibited biphasic phase separation, forming condensates below a critical temperature and dissolving above it **(Figure S1D)**. Notably, the presence of PAR increased the critical temperature **(Figure S1D)**, indicating that PAR-containing FUS condensates require more energy to dissolve.

Building on these simulations, we analyzed the contact frequency between PAR and FUS at a temperature below the critical point, yet high enough to allow efficient sampling of conformational ensembles within tractable computational timescales **(Figure 1G)**. The analysis revealed that PAR preferentially binds to the C-terminal regions of FUS during condensation, particularly the RGG2 domain **(Figure 1G)**.

Supporting this, a construct comprising RGG2-ZnF-RGG3-PY—hereafter referred to as the C-terminal RGG-rich region (CTRR)—restored PAR-binding affinity to levels comparable to full-length FUS and formed condensates with PAR **(Figures 1B–1E, S1B and S1E)**. Therefore, the CTRR of FUS is sufficient to interact and condense with PAR.

### Arginine Residues within RGG Repeats Are Indispensable for Condensation with PAR

The CTRR contains multiple concatenated quasi-repetitive RGG and RG repeats. To test whether these repeats alone are sufficient for PAR-mediated condensation, we titrated increasing concentrations of a synthetic 25-amino acid multivalent polypeptide [RGRGG]_5_^53^. In the presence of 1 µM PAR, significant condensation was observed at ≥ 10 µM [RGRGG]_5_ **(Figures 1H and S1F)**. Conversely, fixing [RGRGG]_5_ at 10 µM revealed that as little as 0.1 µM PAR was sufficient to induce condensation **(Figures 1I and S1G)**. Thus, as with FUS, PAR can drive the condensation of RGG/RG-rich repeats with substoichiometric potency.

We hypothesized that the positively charged arginine residues facilitate condensation via multivalent electrostatic interactions with negatively charged PAR. To test this, we substituted one or both arginine residues in each repeat of the [RGRGG]_5_ peptides with lysine or tyrosine. While both lysine and arginine are positively charged, lysine has a linear side chain terminating in a primary amine, whereas arginine contains a guanidinium with a planar geometry and delocalized electrons **(Figure S1H)**. Tyrosine, although uncharged, is similar to arginine with delocalized electrons in its side chain.

Strikingly, substituting arginine with lysine or tyrosine—partially or fully—abolished condensation under the tested conditions **(Figures 1J and S1I)** and severely diminished PAR-binding affinity **(Figures 1K, S1J and S1K)**, as measured by electrophoretic mobility shift assay (EMSA). These findings underscore the indispensable role of arginine residues—likely due to the distinct chemical properties of the guanidinium groups—in mediating multivalent interactions with PAR to drive condensate formation.

To assess the role of arginine residues in full-length FUS when condensing with PAR, we substituted all 15 arginines in the RGG3-PY region—the minimal region required for FUS–PAR condensation—with alanine **(Figures S1L and S1M)**. This substitution reduced FUS’s overall positive charge, severely impaired condensation, and decreased PAR-binding affinity by 2.7-fold **(Figures 1L, 1M, S1L and S1N)**. Replacing arginine with lysine, which retains overall charge and isoelectric point (pI), also prevented FUS condensation with PAR despite preserving PAR-binding affinity **(Figures 1L, 1M, and S1L–S1N)**.

Fluorescence anisotropy measurements revealed altered hydrodynamic properties for PAR upon binding to FUS **(Figures 1N and S1N)**. Notably, the increase in fluorescence anisotropy was significantly lower when PAR was bound to the R-to-A and R-to-K mutants compared to wild-type FUS **(Figures 1N and S1N)**. These data suggest that, although the R-to-K mutant maintains PAR-binding affinity, it alters the hydrodynamic properties of the FUS–PAR complex, likely disrupting the multivalent interactions required for condensation.

These findings underscore the indispensable role of arginine, extending beyond simple charge complementarity, in mediating proper FUS–PAR interactions required for condensation.

### The Prion-Like Domain (PrLD) of FUS Drives Sustained Growth and Full Condensation Potential of FUS–PAR Assemblies

While CTRR alone was sufficient to form condensates with PAR, these condensates exhibited significantly smaller coverage area than those formed with full-length FUS **(Figures 1D and 2A–2C)**, suggesting that the N-terminal domains further enhance PAR-initiated FUS condensation. Reintroducing only the RGG1 and RRM domains to CTRR (i.e., ΔPrLD) failed to restore the condensation area to the full-length FUS level, indicating that PrLD is essential for achieving full condensation potential **(Figures 2A–2E and S2A)**. Interestingly, removing the PrLD slightly increased FUS’s PAR-binding affinity **(Figures 2F and S2B)**. Therefore, PrLD likely promotes condensation through mechanisms other than enhancing PAR binding.

**Figure 2.**
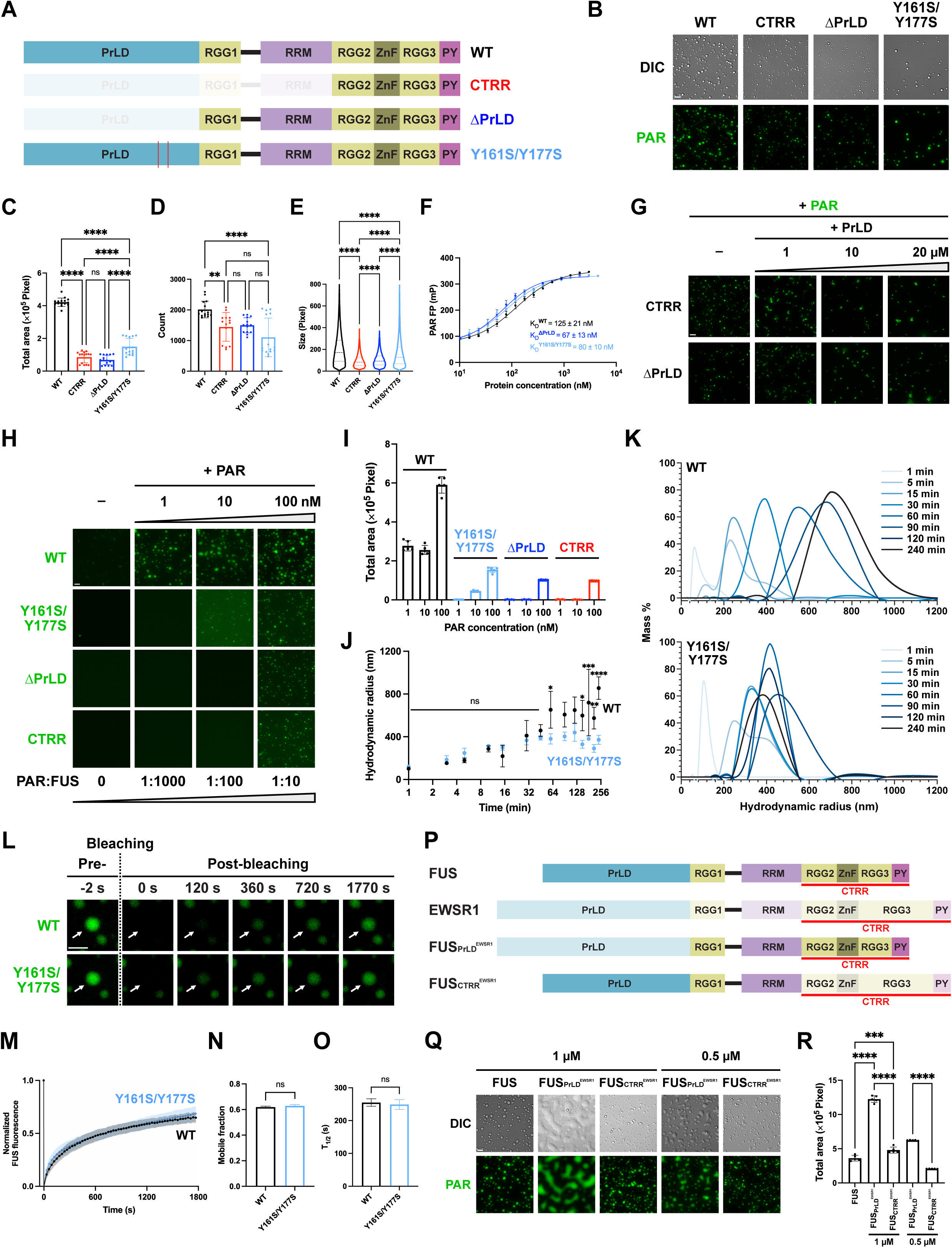
The PrLD of FUS Drives Oligomerization, Enabling Sustained Growth and Maximizing Condensation Potential of FUS–PAR Assemblies. **(A)** Domain structures of CTRR, ΔPrLD, wild-type FUS, and the Y161S/Y177S mutant. **(B)** Representative microscopic images of condensation results for 1 µM of each protein with 100 nM PAR. Fluorescence is from Cy3-labeled PAR. n = 5. Scale bar, 5 µm. **(C-E)** Quantification of the total coverage area of condensates, number of condensates, and condensate size, respectively. For (C) and (D), n = 15; error bars, SD; ordinary one-way ANOVA with Tukey’s multiple comparisons test. For (E), n > 13000; dashed lines in the violin plots represent the 25th, 50th (median), and 75th percentiles; Kruskai-Wallis test with significance levels: not significant (ns), p < 0.05 (*), and p < 0.0001 (****). **(F)** Fluorescence anisotropy assays of wild-type FUS and the ΔPrLD and Y161S/Y177S mutants with 3 nM AF488-PAR. Fluorescence polarization (FP) values of AF488-PAR are plotted and fitted by the Hill equation to determine the K_D_. n = 3; error bars, SD. See also Figure S2B. **(G)** Representative fluorescence microscopic images showing condensation results of 1 µM CTRR or ΔPrLD with increasing concentrations of PrLD in the presence of 100 nM PAR. Fluorescence is from Cy3-labeled PAR. n = 5. Scale bar, 5 µm. See also Figures S2D and S2E. **(H)** Representative fluorescence microscopic images showing condensation results for 1 µM of each protein with a titration of PAR. Fluorescence is from AF488-labeled FUS. n = 5. Scale bar, 5 µm. See also Figure S2G. **(I)** Quantification of the total coverage area of condensates in (H). n = 5; error bars, SD. **(J)** Growth kinetics plot showing dynamic light scattering (DLS) size growth of condensates formed by 1 µM of wild-type FUS or the Y161S/Y177S mutant with 100 nM PAR over 4 h. n = 3; error bars, SD; two-way ANOVA with significance levels indicated at specific time points: not significant (ns), p < 0.05 (*), p < 0.01 (**), p < 0.001 (***), p < 0.0001 (****). See also Figure S2H. **(K)** Representative mass distribution plots of (J) showing percentage mass distribution of condensates by hydrodynamic radius over 4 h. n = 3. See also Figure S2I. **(L)** Representative fluorescence recovery after photobleaching (FRAP) images of condensates formed by 1 µM wild-type FUS or the Y161S/Y177S mutant with 100 nM PAR. Arrows indicate bleached condensates monitored over time. Fluorescence is from the AF488-labeled proteins. n = 5. Scale bar, 5 µm. **(M)** Time-course plot showing normalized fluorescence intensity of FUS in the condensates in FRAP. Fluorescence intensity is normalized to the pre-bleach intensity for each condensate. n = 5; shaded area represents SD. **(N, O)** Comparisons of the mobile fraction and half-time of recovery (T_1/2_) of condensates quantified in FRAP. See **Methods** for the quantification method of mobile fraction and T_1/2_. n = 5; error bars, SD; unpaired t-test with ns represents not significant. **(P)** Domain structures of FUS, EWSR1, and the chimeric proteins FUS_PrLD_^EWSR^^1^ and FUS_CTRR_^EWSR^^1^. FUS_PrLD_^EWSR^^1^ refers to FUS with its PrLD replaced by EWSR1’s PrLD, and FUS_CTRR_^EWSR^^1^ refers to FUS with its CTRR replaced by EWSR1’s CTRR. **(Q)** Representative microscopic images of condensates formed by FUS and FUS chimeric mutants with EWSR1 in the presence of 100 nM PAR. Fluorescence is from Cy3-labeled PAR. n = 5. Scale bar, 5 µm. **(R)** Quantification of the total coverage area of condensates in (P). n = 5; error bars, SD; ordinary one-way ANOVA with Tukey’s multiple comparisons test, with significance levels: p < 0.001 (***) and p < 0.0001 (****).

We hypothesized that PrLD promotes condensation through additional protein-protein interactions. FUS can form homotypic PrLD–PrLD and heterotypic PrLD–RGG intermolecular interactions^42,54–56^. To test whether intermolecular PrLD–RGG interactions drive condensation, we incubated PAR with CTRR or ΔPrLD, which are enriched with RGG repeats, and added increasing concentrations of a PrLD fragment. While these *trans*-complementation conditions slightly altered condensate morphology, they failed to restore condensation to the level observed with the full-length wild-type FUS **(Figures 2G and S2C–S2E)**. Furthermore, similar *trans*-complementation using full-length FUS constructs—which contain an intact PrLD but lack the ability to condense with PAR—also failed to enhance condensation **(Figures S2D and S2F**). These results suggest that intermolecular PrLD–CTRR interactions are not the determinant for full condensation potential.

To assess whether homotypic PrLD–PrLD interactions are crucial for promoting condensation, we constructed a PrLD mutant (Y161S/Y177S), with substitutions previously shown in isolated domains to disrupt self-association *in vitro*^54^. In full-length FUS, these mutations led to a three-fold decrease in the FUS–PAR condensate coverage area, along with significant reductions in condensate number and size **(Figures 2A–2E and S2A)**. Importantly, this reduction was not due to impaired PAR binding **(Figures 2F and S2B)**. Additionally, PrLD mutation (Y161S/Y177S) or removal (ΔPrLD and CTRR) significantly weakened substoichiometric potency when compared to wild-type FUS, with almost no detectable condensation at a 1:1000 PAR:FUS ratio and smaller condensate area at 1:100 **(Figures 2H, 2I, and S2G)**. These results suggest that homotypic PrLD interactions promote the full FUS–PAR condensation potential.

Interestingly, dynamic light scattering (DLS) of the FUS–PAR assemblies revealed that wild-type and Y161S/Y177S mutant FUS assemblies initially exhibited similar growth kinetics; however, while wild-type condensates continued to expand beyond 45 min, mutant assemblies ceased growing **(Figures 2J, 2K, S2H, and S2I)**. This suggests that PrLD interactions are specifically required for the sustained growth phase of FUS–PAR condensation rather than its initial formation. Fluorescence recovery after photobleaching (FRAP) analysis showed comparable mobile fractions and recovery half-times (T_1/2_) for both mutant and wild-type FUS condensates, indicating that PrLD disruption does not impair FUS molecule accessibility for recruitment into condensates **(Figures 2L–2O)**. Together, these findings suggest that PrLD interactions drive oligomerization, enabling sustained growth and maximizing condensation potential of FUS–PAR assemblies.

To investigate the broader role of PrLD as a modular regulator in PAR-mediated condensation, we replaced the PrLD of FUS with the PrLD in EWSR1 (PrLD^EWSR1^) **(Figures 2P and S2J)**, a closely related FET family protein. Compared to FUS’s PrLD, PrLD^EWSR1^ is longer and has additional tyrosine residues. The resulting chimera (FUS ^EWSR1^) exhibited enhanced condensation with PAR compared to the wild-type FUS **(Figures 2Q and 2R)** and displayed hallmark features of spinodal decomposition **(Figures 2Q)**, indicating robust condensation driven by a higher thermodynamic driving force^57^. In contrast, replacing the CTRR of FUS with that of EWSR1 (FUS ^EWSR1^) **(Figures 2P and S2J)**, which is longer and contains more arginine residues than FUS’s CTRR, only slightly increased condensation, and without spinodal decomposition **(Figures 2Q and 2R)**. These findings further highlight the critical—and potentially general—regulatory role of PrLD in promoting PAR-initiated condensation.

### The PAR–CTRR Interaction Opens the FUS Conformation to Promote Condensation

Our findings thus far demonstrate that CTRR is sufficient for PAR-initiated condensation, while PrLD enhances this process. Given that PAR binds directly to CTRR **(Figures 1E and S1E)**, and that PrLD can also interact with RGG repeats enriched in CTRR^42,55^, we hypothesized that PAR disrupts intramolecular PrLD–CTRR interactions in FUS to promote PrLD–PrLD interactions *in trans* for condensation. To test this hypothesis, we first used slab simulations to monitor the distance between PrLD and CTRR within individual FUS molecules upon PAR interaction during condensation. We observed an inverse relationship: as PAR interacts with CTRR and reduces the distance between them, PrLD shifts away from CTRR (increasing distance), leading to a more extended FUS conformation **(Figure 3A and Movie S1)**.

**Figure 3.**
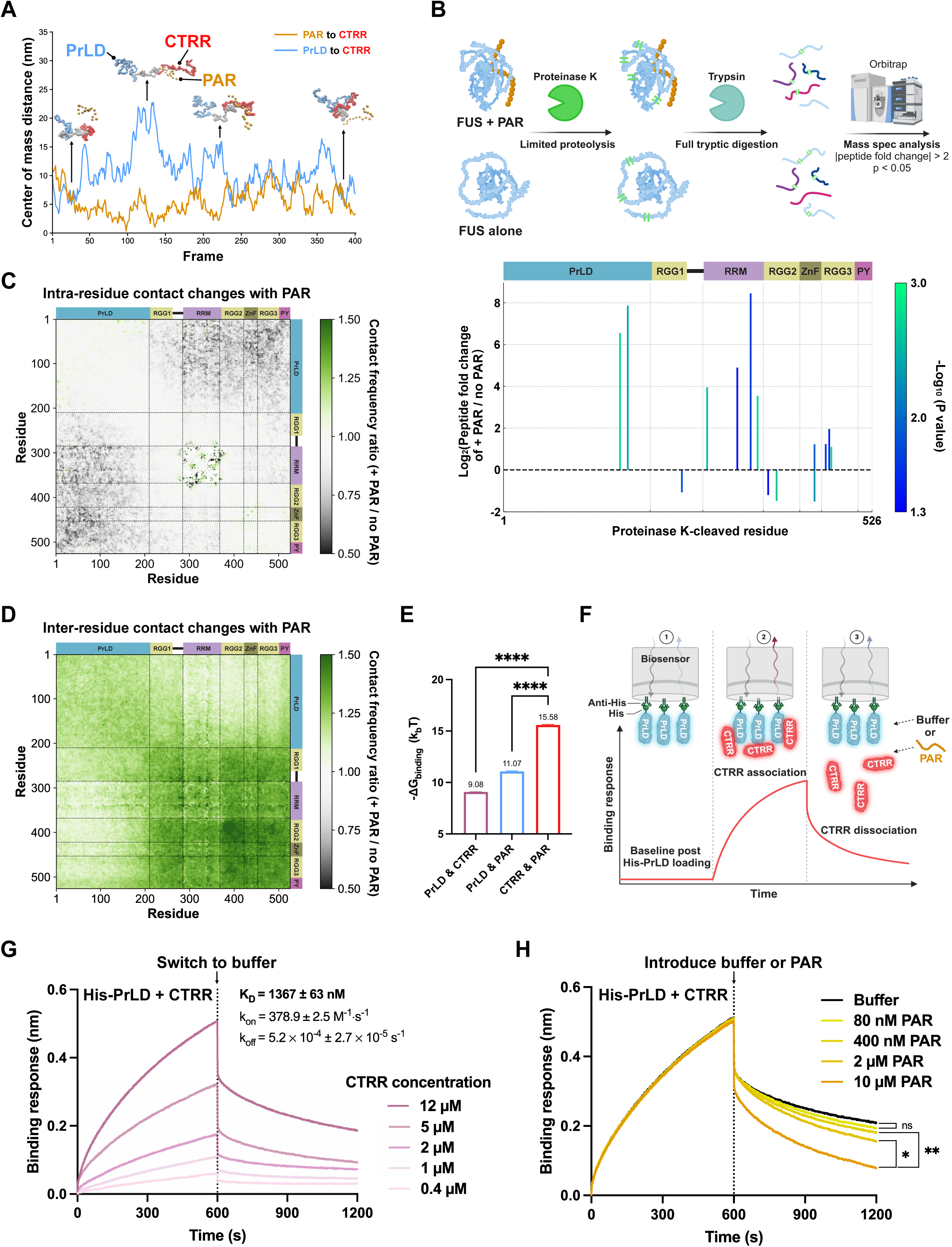
PAR Competes with PrLD Binding to CTRR and Opens FUS Conformation upon Condensation. **(A)** Representative contact map from slab simulations showing the center-of-mass distance between PAR and CTRR, and between PrLD and CTRR across simulation frames. Corresponding structural snapshots on top of the plots highlight the conformations of FUS and the position of PAR at specific frames (See also Movie S1). PrLD is shown in blue, RGG1-RRM in gray, CTRR in red, and PAR in amber. **(B) Upper panel:** Schematic of the limited proteolysis coupled with mass spectrometry (LiP-MS) workflow on FUS with and without PAR. **Lower panel:** LiP-MS accessibility plot showing proteinase K-digested sites across FUS (x-axis) with an absolute value of peptide fold change > 2 in the presence and absence of PAR, and p < 0.05; unpaired Welch’s t-tests; n = 3. The height of each vertical line represents the fold-change, with values above the dashed baseline indicating increased accessibility (more exposed sites) in the presence of PAR, and values below the line indicating decreased accessibility. The color intensity of the vertical lines represents p values, with bright greens indicating more statistically significant changes. **(C, D)** Ratios of intra- (C) and inter- (D) contact frequency changes between FUS residues in the presence and absence of PAR during slab condensation simulations (See **Methods**). Colors represent the ratio of contact frequencies with PAR to those without PAR: gray indicates reduced contact in the presence of PAR, whereas green indicates increased contact in the presence of PAR. The specific color scheme is shown in the legend on the right. The RRM and ZnF regions show noticeable patches in (C) because they are the only two structured domains within FUS. Their relatively fixed secondary and tertiary structures lead to stable intramolecular contact patterns, which persist even during the dynamic condensation process. These structured domains contribute to distinct patterns of intra-residue contact frequency changes compared to disordered regions of FUS. **(E)** The calculated binding free energies for interactions between PrLD, CTRR, and PAR in umbrella sampling simulations (see **Methods**). n = 5; error bars, SD; ordinary one-way ANOVA with Tukey’s multiple comparisons test, p < 0.0001 (****). See also Figure S3A. **(F)** Schematic of biolayer interferometry (BLI) workflow: His-MBP-tagged PrLD is immobilized on biosensors with anti-His, followed by the introduction of CTRR to associate with PrLD. Subsequently, either buffer or PAR is introduced to observe the dissociation of PrLD–CTRR binding. Changes in the wavelength of the reflected light from the biosensor surface indicate association and dissociation responses. **(G)** A representative BLI sensorgram shows measurements of PrLD–CTRR binding at different CTRR concentrations. The association rate (k_on_), dissociation rate (k_off_), and equilibrium dissociation constant (K_D_) are calculated by fitting data to an exponential kinetic binding model (see **Methods**). n = 4. **(H)** Representative BLI sensorgram comparing the effect of buffer and PAR titrations on dissociating PrLD–CTRR interactions. PrLD was immobilized on biosensors and first associated with 12 µM CTRR, followed by introduction of either buffer or PAR during the dissociation phase. n = 3; ordinary one-way ANOVA with Tukey’s multiple comparisons test, with significance levels comparing the projected plateau level at dissociation: not significant (ns), p < 0.05 (*) and p < 0.01 (**). See also Figure S3B.

To experimentally probe this conformational change, we performed limited proteolysis coupled with mass spectrometry (LiP-MS). FUS was incubated with or without PAR for the same duration as in our condensation assays, followed by a brief 10-s digestion with proteinase K, which selectively cleaves structurally accessible regions **(Figure 3B, upper panel)**. Mass spectrometry revealed a significant increase in both the abundance and diversity of proteinase K-digested peptides in the presence of PAR, with pronounced cleavage in the PrLD, RRM, and RGG3 regions **(Figure 3B, lower panel)**. These findings suggest that PAR increases FUS structural accessibility, promoting a more open conformation.

To connect these conformational changes to condensation, we analyzed intra- and intermolecular interactions in slab simulations of 100 FUS molecules, with or without 10 PAR molecules **(Figures 1F, S1C, 3C and 3D)**. In the presence of PAR, intramolecular interactions between PrLD and CTRR decreased **(Figure 3C; cf. Figure 3A and Movie S1** for snapshots), while intermolecular interactions between CTRRs increased **(Figure 3D)**, likely mediated via shared binding to PAR **(Figure 1G)**. This shift suggests that CTRR preferentially binds to PAR over PrLD. Indeed, umbrella sampling simulations (see **Methods**) predicted that the free energy of binding between CTRR and PAR is more favorable than that between CTRR and PrLD **(Figures 3E and S3A)**. Together, these findings support a model in which PAR competitively displaces PrLD from CTRR, promoting a more open FUS conformation.

To experimentally test this competition model, we performed biolayer interferometry (BLI) studies. Briefly, we immobilized PrLD and titrated free CTRR **(Figure 3F-1, 2)**, detecting binding with a K_D_ of 1367 nM **(Figure 3G)**. As expected, the negative control maltose-binding protein (MBP) showed no detectable binding to PrLD **(Figure S3B)**. To determine whether PAR can displace CTRR from PrLD, we pre-bounded CTRR to immobilized PrLD, then introduced either buffer or increasing concentrations of PAR during the dissociation phase **(Figure 3F-3)**. PAR caused significantly greater CTRR dissociation than buffer, in a dose-dependent manner, demonstrating that PAR competes with PrLD for CTRR binding rather than facilitating cooperative interactions among the three components.

In addition, intermolecular interactions between FUS PrLDs are also predicted to increase in the presence of PAR **(Figure 3D)**. Therefore, we propose that FUS condensation is enhanced by a PAR-induced conformational opening of FUS. In this open state, PAR bridges CTRRs across different FUS molecules, while the exposed PrLDs promote homotypic intermolecular interactions that sustain condensation.

### PAR Binding Can Induce an Early Conformational Opening of FUS

While PAR promotes a conformational opening of FUS to enable condensation, we next asked how early these structural changes can occur. Notably, consistent with previous findings^51^, we observed that PAR undergoes compaction upon interacting with FUS prior to visible condensate formation, even at nanomolar concentrations, as indicated by increased FRET efficiency in single-molecule measurements **(Figures 4A–4C and S4A)**. We, therefore, investigated whether FUS conformational opening also occurs at these early, pre-condensation stages.

**Figure 4.**
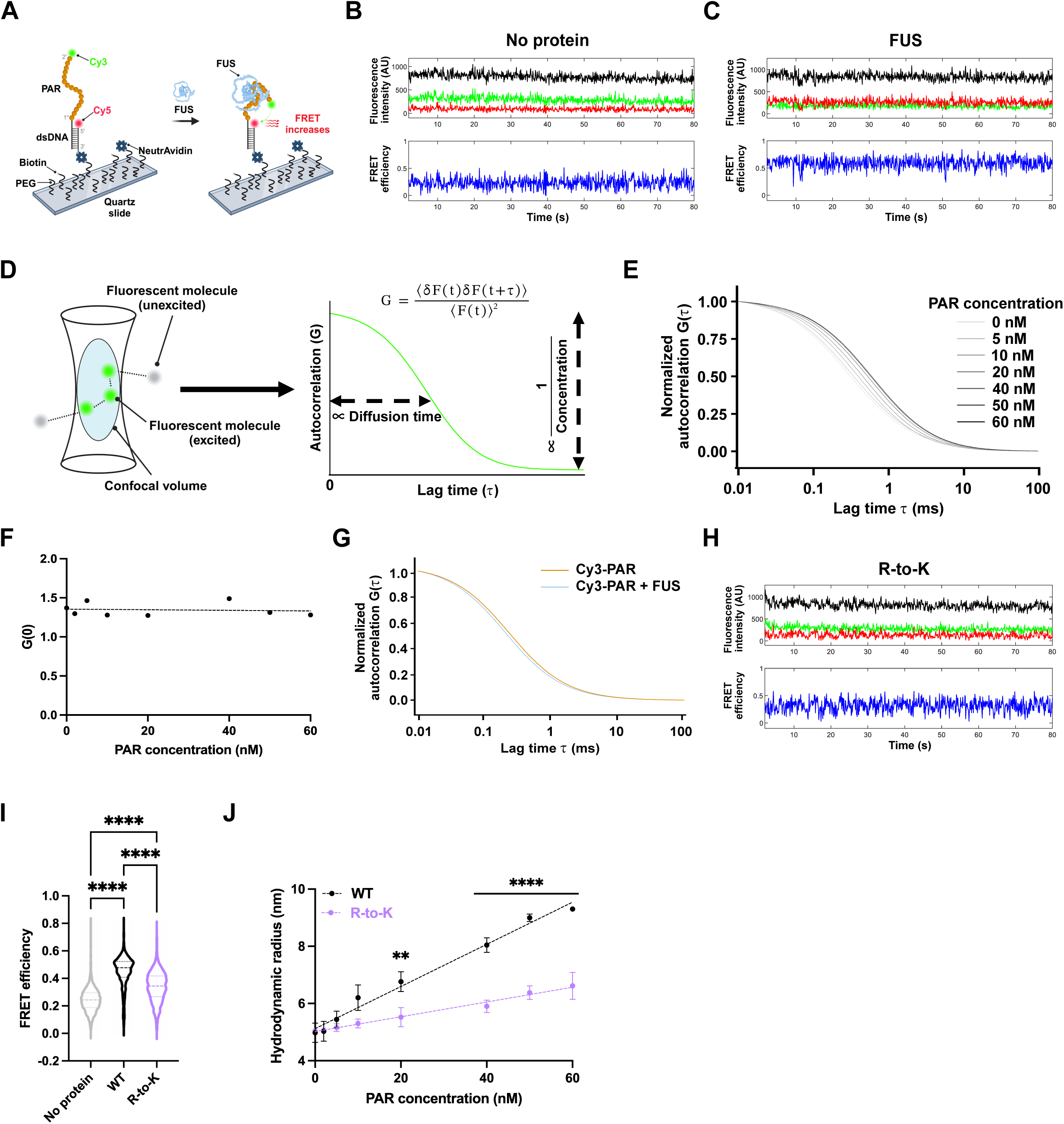
FUS and PAR Undergo Reciprocal Conformational Changes upon CTRR’s Arginine-Mediated Interactions. **(A)** Schematic of single-molecule FRET (smFRET) measurement of the end-to-end distance of a PAR molecule. A partially duplex DNA-PAR molecule was immobilized via NeutrAvidin-biotin interactions onto a PEG-passivated quartz slide, with Cy5 labeled on DNA and Cy3 on PAR. An increased FRET signal indicates PAR compaction upon FUS introduction. **(B, C) Upper panel:** Representative single-molecule traces of Cy3 (green) and Cy5 (red) fluorescence signals, plus total intensity (black, Cy3 + Cy5 intensity offset by 400 for visualization) in response to buffer (no protein) and wild-type FUS (WT, 10 nM), respectively. **Lower panel:** Calculated FRET efficiency corresponding to the upper panel. Frame rate, 10 frames/s. See also (I). **(D)** Schematic of the fluorescence correlation spectroscopy (FCS). **Left:** Fluorescent molecules diffuse in and out of a small confocal volume, where they are excited (green) and emit fluorescence signals. Molecules outside this volume remain unexcited (gray). **Right:** The resulting fluorescence intensity fluctuations are analyzed using the autocorrelation function G(τ) (equation above), which decays over time as molecules diffuse. The characteristic diffusion time, extracted from the decay curve, reflects the hydrodynamic radius of the molecules. The autocorrelation amplitude at τ = 0, G(0), is inversely proportional to the number of fluorescent molecules and thus to their concentration. **(E)** Representative FCS autocorrelation curves showing a dose-dependent increase in diffusion time (rightward shift) of Cy3-labeled FUS (20 nM) in response to increasing concentrations of unlabeled PAR (0–60 nM). n = 3. See also (J). **(F)** The initial autocorrelation amplitude G(0) is plotted against increasing PAR concentrations. The linear regression fitting line is shown as a dashed line. No significant correlation is observed between G(0) and PAR concentrations, indicating that higher-order FUS assemblies did not form. See also (D). **(G)** Representative FCS autocorrelation curves showing the diffusion time of Cy3-labeled PAR (60 nM) in the presence (blue) and absence (amber) of unlabeled FUS (20 nM). **(H)** Representative single-molecule traces and calculated FRET efficiency of PAR interacting with the R-to-K mutant (10 nM). Frame rate, 10 frames/s. See also (I). **(I)** Violin plots of PAR FRET efficiencies in response to 10 nM protein. n > 800; dashed lines represent the 75th percentile, the median (50th percentile), and the 25th percentile; Kruskal-Wallis test, p < 0.0001 (****). **(J)** Hydrodynamic radius of wild-type FUS and the R-to-K mutant in response to increasing PAR concentrations, measured by FCS. Protein concentration: 20 nM. n = 3; error bars represent SD. Statistical analysis was performed using two-way ANOVA, with significance levels: p < 0.01 (**) and p < 0.0001 (****).

To assess how FUS conformation changes upon PAR interaction at nanomolar concentrations, we used fluorescence correlation spectroscopy (FCS) to monitor the diffusion of Cy3-labeled FUS and infer changes in its hydrodynamic radius **(Figure 4D)**. Increasing concentrations of PAR led to a dose-dependent rightward shift in FUS autocorrelation curves **(Figure 4E)**, consistent with conformational expansion. While this shift supports the idea of PAR-induced FUS opening, alternative explanations— such as FUS oligomerization or stable FUS–PAR complex formation—could also account for the observed change.

To test for FUS oligomerization, we analyzed the initial amplitude of the FCS autocorrelation function, G(0), which is inversely proportional to the number of diffusing fluorescent molecules **(Figure 4D, right)**. G(0) remained nearly constant across the tested PAR concentration range **(Figure 4F)**, arguing against oligomerization.

Next, we tested whether a stable FUS–PAR complex forms by swapping the dye and performing FCS on Cy3-labeled PAR at its highest concentration used during the Cy3-FUS experiments (see **Figure 4E)**. If a stable complex were to form, we would expect a rightward shift in the PAR autocorrelation curve upon adding FUS due to an increased hydrodynamic radius. However, no such shift was observed **(Figure 4G)**. Instead, we detected a slight leftward shift, consistent with PAR compaction as suggested by the FRET experiments **(Figures 4A–4C and S4A)**. Furthermore, as in prior studies^4^, our single-molecule experiments confirmed that FUS–PAR interactions at these concentrations are dynamic and transient **(Figures S4B and S4C)**. Therefore, we infer that PAR binding can induce an early conformational opening of FUS, even at sub-condensation concentrations, without requiring oligomerization or stable complex formation.

### Arginines within CTRR Are Required for Reciprocal Conformational Changes in FUS and PAR

Our data suggest that PAR exhibits altered hydrodynamic properties when interacting with FUS through lysine rather than arginine in the CTRR of FUS, despite similar binding affinities (see WT vs. R-to-K mutant; **Figures 1L–1N, and S1L–S1N)**. Consistently, FRET experiments showed that the R-to-K mutant induced significantly less PAR compaction **(Figures 4H, 4I, and S4A)**, highlighting the unique role of arginine residues within FUS’s CTRR in mediating PAR structural dynamics during interaction.

To test whether these arginine residues are required for PAR-induced FUS conformational opening, we used FCS to compare the hydrodynamic radii of Cy3-labeled wild-type and R-to-K mutant FUS in response to increasing PAR concentrations. The R-to-K mutant exhibited a significantly reduced expansion in hydrodynamic radius compared to wild-type FUS **(Figure 4J)**. At 60 nM PAR, the R-to-K mutant’s hydrodynamic radius was 40% smaller than that of wild-type FUS **(Figure 4J)**, indicating compromised PAR-induced conformational opening. These findings suggest that specific arginine-mediated interactions are critical for the induction of a conformationally responsive FUS state. This behavior is consistent with a nucleation-competent intermediate^44^, in which transient PAR engagement promotes a conformational switch that primes FUS for sustained condensation.

Taken together, these results demonstrate that FUS and PAR undergo reciprocal conformational changes upon CTRR’s arginine-mediated interactions. These findings underscore the unique role of arginine, which cannot be functionally substituted by lysine, in driving the FUS–PAR interactions required to initiate condensation.

### PrLD Interactions Sustain PAR-Initiated FUS Condensation upon PAR Degradation *in vitro*

As a transient trigger in DNA damage response, PAR synthesized by PARP1 is rapidly degraded by its glycohydrolase PARG, transitioning the repair process to a PAR-independent phase^27^. Yet, FUS persists at repair foci well beyond PAR degradation^22,32,38^. Similarly, *in vitro*, PAR-initiated FUS condensation persists even after PAR degradation by PARG **(Figure 5A)**^4^, whereas no condensates formed under the same conditions when FUS was initially incubated without PAR **(Figure S5A)**. These findings suggest that PAR not only initiates FUS condensation but also induces a state in which FUS condensates persist for a sustained period even after PAR is degraded, likely through biophysical changes within FUS.

**Figure 5.**
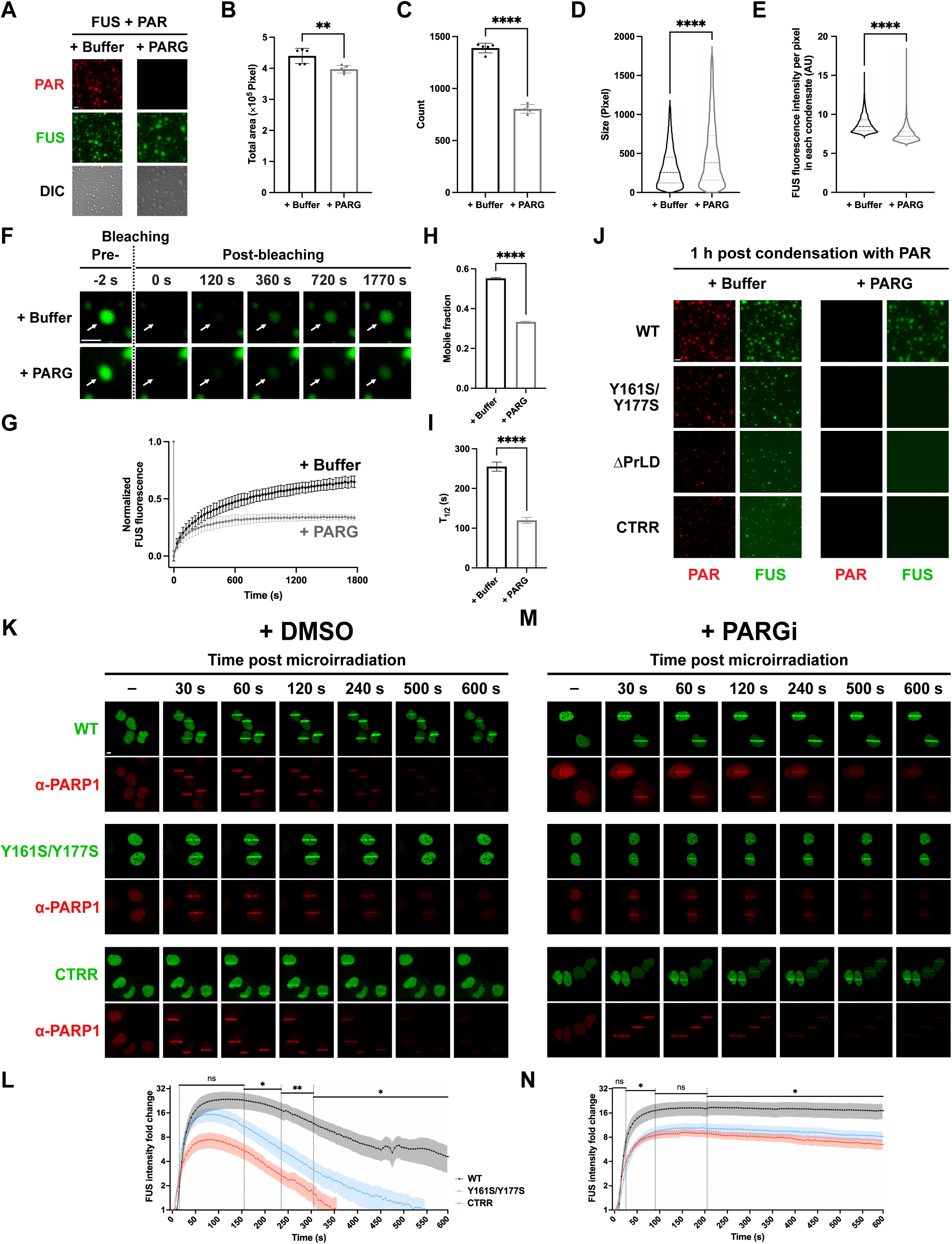
PrLD Interactions Sustain PAR-Initiated FUS Condensation upon PARG-Mediated PAR Degradation in DNA Damage Response. **(A)** Representative microscopic images of condensates formed by 1 µM FUS (green) and 100 nM PAR (red) after 1 h, followed by treatment with either buffer or PARG for an additional 1 h. n = 5. Scale bar, 5 µm. **(B, C)** Bar graph comparisons of the total coverage area of condensates and the number of condensates with and without PARG treatment. n = 5; unpaired t-test, p < 0.01 (**) and p < 0.0001 (****). **(D, E)** Violin plot comparisons of condensate size and FUS fluorescence intensity per pixel within each condensate, with and without PARG treatment. n > 4000; dashed lines represent the 75th percentile, the median (50th percentile), and the 25th percentile; Mann-Whitney test, p < 0.0001 (****). **(F)** Representative FRAP images of FUS–PAR condensates with or without PARG treatment. Arrows indicate bleached condensates monitored over time. Fluorescence is from AF488-labeled proteins. N = 5. Scale bar, 5 µm. **(G)** Time-course plot showing normalized fluorescence intensity of FUS in the condensates from FRAP in (F). Fluorescence intensity is normalized to the pre-bleach intensity for each condensate. n = 5; error bars, SD. **(H, I)** Comparisons of the mobile fraction and T_1/2_ of condensates quantified in FRAP. The quantification method for mobile fraction and T_1/2_ is described in the **Methods**. n = 5; error bars, SD; unpaired t-test, p < 0.0001 (****). **(J)** Representative microscopic images of condensates formed by 1 µM of wild-type FUS or corresponding mutants (green) with 100 nM PAR (red) after 1 h, followed by treatment with either buffer or PARG for an additional 1 h. The protein-only controls are listed on the right. n = 5. Scale bar, 5 µm. See also Figure S5B. **(K–N)** Laser strip assays in U2OS cells transiently transfected with EGFP-FUS (wild-type, Y161S/Y177S, or CTRR; green) with or without 1 µM PARG inhibitor (PARGi; PDD 00017273) pretreatment, and corresponding FUS enrichment intensity quantification. (K, M) Representative time-course images of EGFP-FUS localization at DNA damage sites with DMSO (K) or PARGi (M) pretreatment. Endogenous PARP1 (red) was detected using an RFP-tagged anti-PARP1 chromobody (α-PARP1). See also Figure S5E. (L, N) Normalized fold change of FUS enrichment intensity, calculated as EGFP signal at DNA damage lines relative to pre-microirradiation intensity values of the same regions later being quantified as DNA damage lines, corresponding to (K) and (M). n ≥ 15; shaded areas represent SD. Statistical significance between wild-type FUS and the Y161S/Y177S mutant was assessed using multiple unpaired Welch’s t-tests: not significant (ns), p < 0.05 (*) and p < 0.01 (**). Dashed vertical lines indicate ranges with consistent significance levels.

Indeed, following PARG treatment, we observed fewer yet larger condensates, accompanied by a slight decrease in the total condensate area and lower FUS concentration within these condensates **(Figures 5B–5E)**. FRAP analysis further revealed a decreased mobile fraction of FUS within the condensates **(Figures 5F–5I)**. Collectively, these changes likely indicate that PAR-mediated interactions are disrupted, while residual intermolecular FUS interactions sustain the condensates.

To identify the molecular determinants that sustain FUS condensates following PAR degradation, we examined various condensate-forming mutants for their PARG sensitivity *in vitro*. Unlike wild-type FUS, condensates formed by the PrLD-deficient mutant Y161S/Y177S, as well as by ΔPrLD and CTRR—both of which lack the entire PrLD—dissipated rapidly following PARG treatment **(Figures 5J and S5B)**. These findings strongly suggest that PrLD, and likely its intermolecular interactions, are essential for sustaining PAR-initiated condensation following PAR degradation.

### PrLD Interactions Sustain PAR-Initiated FUS Condensation upon PAR Degradation in Cells

Previous studies demonstrated that PAR-dependent FUS enrichment appears as condensed, liquid-demixing stripes around DNA damage sites induced by laser microirradiation in cells^22,33^. To investigate whether the molecular determinants identified *in vitro* are relevant in cells, we induced DNA damage in the nuclei of U2OS cells transfected with EGFP-FUS or its mutants **(Figure S5C)**. An RFP-tagged anti-PARP1 chromobody, which enables real-time tracking of endogenous PARP1^58^, showed that PARP1 was recruited to damage sites within 5 s **(Figure S5D)**, consistent with previous findings^59^. Wild-type FUS became enriched around DNA damage sites within 30 s, reaching peak fluorescence intensity between 60 and 120 s^31,32^ **(Figure 5K, upper panel)**. Although FUS gradually dissipated thereafter, its enrichment persisted for the full 10-min observation period^22,32,37,38^ **(Figures 5K, top panel, and S5E)**.

In contrast, the Y161S/Y177S mutant and CTRR were enriched around damage sites but with significantly lower fluorescence intensity, mirroring their reduced potency in PAR-induced condensation observed *in vitro*. Furthermore, these mutants dissipated more rapidly, becoming undetectable by ∼450 s **(Figures 5K, 5L, and S5E)**. Notably, nuclear FUS intensity prior to microirradiation remained comparable between wild-type and mutant-expressing cells **(Figure S5F)**. Thus, differences in FUS enrichment duration are not due to variations in FUS expression levels but rather stem from the absence of a functional PrLD, underscoring its critical role in sustaining FUS enrichment at DNA damage sites. This persistence in cells further supports the notion that PAR triggers a conformational transition in FUS that is actively maintained through PrLD-mediated interactions, effectively decoupling the persistence of the condensate from its initiating signal.

Our *in vitro* data indicated that condensates formed by the Y161S/Y177S mutant and CTRR are more susceptible to dissipation upon PARG treatment than wild-type FUS condensates **(Figures 5J and S5B)**. We hypothesized that the differences in FUS enrichment duration in cells result from their distinct responses to PAR degradation by endogenous PARG: Wild-type FUS sustains enrichment longer after PAR degradation, whereas mutants dissipate more rapidly. If true, inhibiting PARG activity should prolong FUS enrichment and equalize persistence between wild-type and mutant proteins. To test this, we treated cells with the PARG inhibitor PDD 00017273 (PARGi)^60^. PARG inhibition did not significantly alter nuclear FUS protein levels **(Figure S5F)**, but it markedly prolonged FUS enrichment. Importantly, both wild-type and mutant FUS exhibited comparable enrichment persistence while maintaining their distinct enrichment levels **(Figures 5M, 5N, and S5E)**. These results support our hypothesis that the rapid dissipation of mutants stems from their inability to sustain enrichment after PAR degradation—a function dependent on an intact PrLD.

Together, our findings indicate that while PAR initiates FUS condensation, PrLD-mediated interactions are critical for sustaining condensation after PAR degradation, both *in vitro* and in cells.

## DISCUSSION

### PAR-Induced Conformational Changes Are Critical for Condensation Hysteresis

While nucleic acids commonly act as compositional structural scaffolds in biomolecular condensates^13,14^, the noncanonical nucleic acid PAR serves as a transient trigger in DNA repair foci. Unlike DNA or RNA, PAR contains an additional phosphate group per monomer and two flanking ribose moieties, one carrying an adenine base capable of π-stacking. These features may confer a unique electrostatic and spatial configuration that enhances PAR’s potency in driving protein condensation^61^. While high concentrations of nuclear RNA typically maintain FUS in a soluble state^62^, PAR synthesized upon DNA damage relocalizes FUS to these damage sites and promotes the formation of FUS-enriched repair foci, positioning PAR as a spatiotemporal inducer of biomolecular condensation.

PAR is essential for initiating FUS condensation in DNA repair, but the condensates persist even after PAR is degraded. Both experimental data and molecular dynamics simulations support a bi-modular mechanism in which FUS’s two intrinsically disordered termini, CTRR and PrLD, coordinate spatially to shift the condensation from a PAR-driven to a protein-driven state. Specifically, PAR binding to CTRR triggers conformational opening that displaces intramolecular contacts between CTRR and PrLD. Condensation occurs at this stage most likely through multivalent interactions between PAR and CTRRs from multiple FUS molecules. The exposed PrLD enables interactions *in trans* that maintain condensation after PAR is lost **(Figure 6)**. These PAR-induced conformational changes—and the resulting persistence of condensates— may be made possible due to the conformational flexibility inherent to these two intrinsically disordered modules.

**Figure 6.**
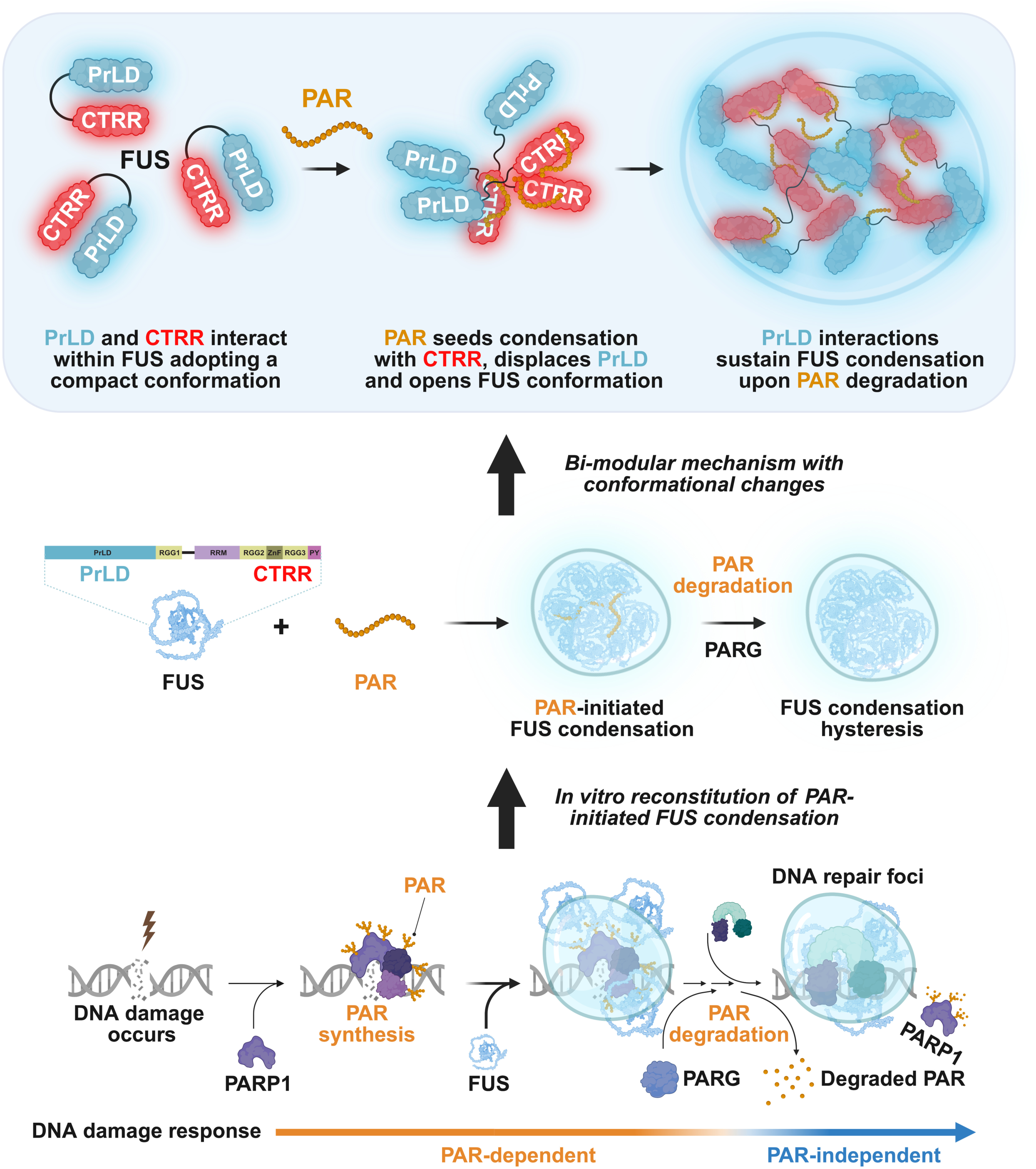
A Bi-Modular Model of How PAR Initiates and Primes Sustained Condensation of FUS

A similar mechanism has been observed for G3BP1, where RNA binding induces a conformational switch that enables condensation and stress granule assembly^20,63,64^. However, unlike G3BP1, FUS condensates persist even after the initiating nucleic acid is removed. This difference may stem from the transient and dynamic nature of FUS– PAR interactions combined with the prion-like character of FUS’s oligomerization domain, which can adopt multiple conformations^65,66^—some of which may stabilize condensates independent of PAR. Supporting this model, our single-molecule studies show that even nanomolar concentrations of PAR are sufficient to induce FUS conformational opening via transient interactions **(Figures 4D–4G, S4B and S4C)**, suggesting that a transient trigger can impose a lasting effect on the condensate state.

This behavior is reminiscent of classical prions, which undergo a transition into a persistent, self-sustaining conformational state following a transient nucleation event^44^. In this context, PAR may act as a structural trigger, converting FUS into a metastable conformation stabilized through PrLD–PrLD interactions. These findings suggest that conformational memory—long considered a hallmark of pathological prion-like aggregation—may represent a broader principle governing the regulated persistence of biomolecular condensates.

### PrLD Is Dispensable for Initiation but Required for FUS Condensation Hysteresis

Although the PrLD has long been considered the primary driver of protein condensation^67,68^, our data reveal that it is neither necessary nor sufficient for PAR-induced FUS condensation. The CTRR alone can nucleate condensate formation in response to PAR. However, the PrLD is indispensable for converting these PAR-induced condensates into a sustained, protein-driven state, which is particularly relevant in DNA repair, where PAR is rapidly synthesized and degraded. This finding aligns with an artificial system showing that light-induced FUS PrLD condensates can retain spatial memory of transient light patterns in cells^69^.

Moreover, the FUS PrLD interacts with the PrLDs of other RNA-binding proteins such as RBM3, CPEB2, TIA1, hnRNPA1, and hnRNPA2^39,56,70^. The PrLD of hnRNPA2, in turn, can interact with those of CIRBP, RBM3, and hnRNPA1, building a broader network of PrLD–PrLD interactions^71^. While many of these proteins participate in DNA repair^38,72^, they cannot condense independently at physiological concentrations^42,73,74^. Thus, PrLD-mediated interactions may not only stabilize the initial PAR-triggered FUS condensate but also facilitate the recruitment of additional PrLD-containing factors, providing a mechanistic framework to explore the maturation and dynamic compositional remodeling of DNA repair foci.

Interestingly, similar nucleic acid-triggered hysteresis is observed in several innate immune sensors, which form oligomeric filaments that remain stable even after the activating DNA trigger has degraded^5,6^. This suggests a broader principle in which nucleic acid-triggered condensation transitions into a self-sustaining state via oligomerization domains. This model may apply to diverse cellular processes and warrants further investigation.

### Modeling the Co-Condensation of Full-Length FUS and PAR Homopolymer

One key advance of our study is the use of molecular dynamics simulations to elucidate how a noncanonical nucleic acid, PAR, induces protein condensation. To capture PAR’s physical behavior and interactions with FUS, we developed a customized coarse-grained model of PAR, optimized using smFRET data. This model enabled the simulation of PAR’s structure and dynamics during condensation, which has been a unique challenge due to the limited structural knowledge of PAR. In terms of protein modeling, while previous work has primarily focused on isolated FUS domains such as the PrLD and RGG1 motif^50,55,75^, we extended this by modeling full-length FUS, incorporating both its structured and intrinsically disordered regions^76,77^. Our simulations revealed key molecular contacts and demonstrated that PAR binding induces a conformational opening of FUS, which facilitates intermolecular interactions and drives sustained condensate growth. Together with experimental validation, these findings offer a foundational framework for understanding how PAR, present in diverse condensates^15^, regulates protein assembly and dynamics with unique structural features. Extending this approach to other nucleic acid–triggered condensates may yield novel mechanistic insights—particularly into conformational dynamics^78^.

### Disease Implications of Bi-Modular Control of Nucleic Acid-Induced Condensation Hysteresis

The bi-modular mechanism by which transient signals like PAR initiate long-lived biomolecular condensates holds significant implications for human disease. A key feature of this mechanism is the nucleic acid–sensing module exemplified by the intrinsically disordered FUS’s CTRR, where arginine-mediated interactions play an indispensable role in PAR-induced conformational transition and condensation. This phenomenon is reminiscent of the behavior of toxic arginine-rich dipeptide repeat proteins (R-DPRs) produced by *C9orf72* repeat expansions—the most common genetic cause of ALS and FTD^79^. In this context, PAR not only induces the condensation of R-DPRs but also enhances their toxicity^80^, highlighting a disease-relevant role for PAR– arginine interactions in neurodegeneration.

Many other proteins also exhibit a similar bi-modular architecture as FUS. Among the ∼240 human proteins with PrLDs, 80 bind DNA, 72 bind RNA, and 78 bind PAR^81–85^— suggesting that this bi-modular mechanism could extend beyond FUS. Whether other PrLD-containing proteins lacking RGG-rich regions—such as TDP-43 or hnRNPA1^67,71^—exhibit similar bipartite behavior remains unclear. However, since both TDP-43 and hnRNPA1 can condense with PAR^73,86^, it is plausible that alternative PAR-binding regions may substitute for the role attributed to FUS’s CTRR.

Notably, like FUS^67^, many of these PAR-binding proteins—including TDP-43, hnRNPA1, R-DPRs, and α-synuclein—are implicated in neurodegenerative diseases^87–89^, where they often form aberrant, persistent inclusions or assemblies^88,90^. Furthermore, PAR levels are often elevated in neurodegeneration due to chronic DNA damage and PARP1 hyperactivation^90–93^. Our previous work demonstrated that, unlike liquid-like droplets, high concentrations and longer lengths of PAR can drive FUS to form solid-like aggregates^4^. Similarly, PAR binds to α-synuclein and promotes the conversion of Parkinson’s disease–related pathological α-synuclein into more misfolded, toxic oligomers both *in vitro* and *in vivo*, accelerating its pathological transmission^90^. These findings suggest that PAR generated during cellular stress may trigger pathological condensation events that evolve into self-sustaining, irreversible prion-like aggregates.

This principle is further supported by our finding that replacing the PrLD of FUS with that of EWSR1 alters PAR-mediated condensate morphology—from discrete droplets to a spinodal, wave-like pattern. This domain-level modularity also mirrors the architecture of oncogenic fusion proteins in cancer, where PrLDs are frequently fused to nucleic acid-binding modules of transcription factors^94^. Such fusions drive condensate formation at specific genomic loci, leading to dysregulated gene expression and oncogenesis^95,96^.

For instance, FUS-DDIT3 in myxoid liposarcoma and EWSR1-FLI1 in Ewing sarcoma each combine a PrLD with a DNA-binding domain to form transcriptionally active nuclear condensates^97^. *In vitro*, DNA enhances the fiber formation of the FUS-FLI1 fusion protein^98^, reinforcing the idea that nucleic acid binding can nucleate and stabilize disease-relevant assemblies.

Together, these findings suggest that nucleic acid–triggered, PrLD-mediated condensation is a common theme in disease, and that individual modules within this bi-modular framework may represent distinct therapeutic targets. Elucidating the mechanisms that govern regulated condensation hysteresis—and determining whether it can transition into irreversible pathological aggregation—may provide critical insights for developing treatments in neurodegeneration and cancer.

### Limitations

Our approach combines *in vitro* bulk, single-molecule biochemical, biophysical, structural mass spectrometry, *in silico* simulations, and cellular techniques. While these methods are powerful for dissecting specific molecular mechanistic details, they may not fully capture the complexity of the cellular environment. Additional factors or modifications in cells could influence FUS–PAR dynamics, potentially adding layers of spatial regulatory organization to DNA repair foci. Additionally, PAR can exist in diverse forms, including free polymers^99^ and conjugated with proteins such as PARP1 and histones. PAR can also vary in length and exhibit branching^100^. Understanding how these various forms of PAR influence condensation and DNA repair foci would be highly valuable. Together, these insights could deepen our understanding of PAR-mediated condensation.

## RESOURCE AVAILABILITY

### Lead Contact

Further information and requests for resources, reagents, or data should be directed to and will be fulfilled by the lead contact, Anthony K. L. Leung.

### Materials Availability

All unique reagents generated in this study (e.g., FUS mutants, PAR probes, etc.) are available from the lead contact upon request without restriction.

### Data and Code Availability

All data supporting the findings of this study are available from the lead contact upon reasonable request. This study used previously published analysis scripts for smFRET and single-molecule tracking (see **Methods**) and does not report any new custom code. Any additional information required to reanalyze the data is available from the lead contact upon request. The smFRET data acquisition and analysis package is freely available at https://github.com/Ha-SingleMoleculeLab. IDL software can be accessed at https://www.exelisvis.co.uk/ProductsServices/IDL.aspx, and MATLAB is available at https://www.mathworks.com/; both require academic or individual licenses obtainable from their respective providers.

## METHODS

### Purification of FUS Protein

Recombinant FUS protein and its variants were expressed and purified using a multistep protocol. Competent *E. coli* BL21 (DE3 Rosetta 2) cells were transformed with the His-MBP-FUS plasmid via heat shock and selected on kanamycin-supplemented agar plates. Starter cultures in LB medium were used to inoculate Terrific Broth (TB) containing kanamycin, and protein expression was induced with 0.2 mM IPTG when the optical density at 600 nm (OD600) reached 1. Cultures were incubated at 16 °C for 16 h and harvested by centrifugation at 4,000 × g for 30 min at 4 °C. Bacterial pellets were stored at -80 °C until lysis.

Cell lysis was performed in a buffer containing 25 mM HEPES (pH 7.4), 500 mM NaCl, 20% (v/v) glycerol, 4 mM DTT, 1% (v/v) IGEPAL (NP-40), 2 mg/mL lysozyme, 0.4 mM phenylmethylsulfonyl fluoride (PMSF), 400 mM L-Arginine monohydrochloride, 0.08 tablets/mL cOmplete EDTA-free protease inhibitors, and 50 µg/mL RNase A. Cells were disrupted by sonication (55% amplitude, 8-s on/off pulses, for a total of 6 min) on ice, and lysates were clarified by ultracentrifugation at 25,000 × g for 30 min. The resulting supernatant was filtered through a 0.2 µm filter to remove particulates.

The clarified lysate was subjected to amylose affinity purification using amylose resin (New England Biolabs, E8021S) to capture His-MBP-tagged FUS. Proteins were eluted in a buffer containing 50 mM HEPES (pH 7.4), 500 mM NaCl, 20% (v/v) glycerol, 2 mM DTT, and 20 mM maltose monohydrate. Eluates were diluted into a low-salt buffer containing 100 mM NaCl, and filtered through a 0.2 µm membrane. Further purification was performed using a 5 mL HiTrap Heparin HP affinity column (Cytiva, 17040701) with stepwise elution at increasing NaCl concentrations (10%, 20%, 50%, and 100% Buffer B, where Buffer B contained 1 M NaCl) on a Bio-Rad NGC FPLC system. The final fractions containing FUS were pooled and exchanged into a storage buffer containing 50 mM HEPES (pH 7.5), 500 mM NaCl, 20% (v/v) glycerol, and 2 mM DTT. The protein was concentrated using a centrifugal filter with a 50 kDa molecular weight cutoff, aliquoted, flash-frozen, and stored at -80 °C. The purity and integrity of the protein were assessed at each step by SDS-PAGE with Coomassie Brilliant Blue staining. Protein concentrations were determined using a Thermo Scientific NanoDrop One Microvolume spectrophotometer at 280 nm.

### Purification of TEV Protease

The recombinant Tobacco Etch Virus (TEV) protease (S219V mutant) was expressed in *E. coli* BL21 (DE3) cells transformed with the pRK793 plasmid (Addgene #8827).

Protein expression was induced with 1 mM IPTG at an OD600 of 1.0, followed by overnight incubation at 16 °C. Harvested cells were resuspended in lysis buffer containing 20 mM HEPES (pH 8.0), 500 mM NaCl, 10% (v/v) glycerol, 25 mM imidazole, 2 mM DTT, 2 mg/mL lysozyme, 0.4 mM PMSF, 400 mM L-Arginine monohydrochloride, 0.08 tablets/mL cOmplete EDTA-free protease inhibitors, and 50 µg/mL RNase A. The cells were lysed by sonication and clarified by centrifugation. The lysate was purified using a HisTrap HP column (Cytiva, 17524802) with gradient imidazole elution (25–500 mM) and further purified on a HiLoad 16/600 Superdex 75 pg size exclusion column (Cytiva, 28989333) with SEC buffer (20 mM HEPES, 500 mM NaCl, 10% (v/v) glycerol, 1 mM DTT) using a Bio-Rad NGC FPLC system. The purified protein was concentrated, aliquoted, flash-frozen, and stored at -80 °C.

### Fluorescent Labeling of FUS Protein

To label FUS protein with Alexa Fluor 488 (AF488), 16 µM FUS protein was mixed with 160 µM AF488-NHS ester (Lumiprobe, 11820) in the presence of 0.1 M sodium bicarbonate (pH 8.3) in a final reaction volume of 550 µL. The reaction was conducted in 1.5 mL Pierce low protein binding microcentrifuge tubes (Thermo Scientific, 90411), protected from light, and incubated at 23 °C for 45 min at 800 rpm in an Eppendorf thermomixer. Excess dye and contaminants were removed using a 5 mL HiTrap Desalting column (Cytiva, 17140801) on a Bio-Rad NGC FPLC system equilibrated with the FUS storage buffer (50 mM HEPES (pH 7.5), 500 mM NaCl, 20% (v/v) glycerol, and 2 mM DTT, and filtered through a 0.45 µm membrane prior to use). The labeled sample was filtered through a 0.45 µm membrane, injected into a 500 µL sample loop, and passed through the column at a flow rate of 1 mL/min. Peak fractions (∼2 mL) were collected, and protein and dye concentrations were quantified using a Thermo Scientific NanoDrop One Microvolume UV-Vis Spectrophotometer at 280 nm and 488 nm, respectively. Labeling efficiency was determined to be 60–70% on average, and the labeled protein was concentrated, aliquoted, flash-frozen, and stored at -80 °C. Labeling quality was evaluated by SDS-PAGE, followed by fluorescence imaging using a Li-Cor scanner and Coomassie Brilliant Blue staining to confirm protein integrity.

### Fluorescent Labeling of PAR

The 16-mer PAR was synthesized and purified previously^4,51^. Fluorescent labeling of PAR was performed using ELTA^101^. Reactions (100 µL) contained 800 pmol PAR, fluorescent dATP (40 µM), ELTA buffer (20 mM Tris-HCl, pH 7.5, 20 mM Mg(OAc)_2_, 2.5 mM DTT), and OAS1 with poly(I:C), and were incubated at 37 °C for 2 h. After incubation, poly(I:C) was digested with RNase R. Labeled PAR was purified by ethanol precipitation, lyophilized, and resuspended in water. Purity was confirmed by gel electrophoresis, and concentrations were quantified by UV-Vis spectrophotometry. Samples were stored at -80 °C.

### *In Vitro* Reconstitution of Condensates and Microscopic Analysis

Condensate reconstitution assays were performed *in vitro* to investigate the formation of protein condensates under various conditions with PAR. Reactions were conducted in μ-Plate Angiogenesis 96-well ibitreat chamber slides (Ibidi, 81506), with each well pre-incubated with 50 µL of 100 mg/mL bovine serum albumin (BSA) to minimize nonspecific adhesion. Reaction mixtures (40 µL) were prepared in a buffer containing 50 mM Tris-HCl (pH 7.5), 100 mM NaCl, 1 mM DTT, 1 mM EDTA, and 2% (v/v) glycerol. For fluorescence experiments, 10% of the protein and/or PAR were fluorescently labeled, with the remaining 90% unlabeled. Condensation was initiated by adding TEV protease at a final concentration of 4 µM to remove the His-MBP tag from the protein. Samples were gently mixed and incubated in 0.5 mL DNA LoBind Tubes (Eppendorf, 022431005) at 23 °C for 2 h to allow condensation. If PARG treatment was applied, previously purified recombinant PARG^4^ was added to the condensation reaction at a final concentration of 10 µM after 1 h of incubation, followed by an additional hour to allow PAR degradation. After the incubation period, samples were transferred to the pre-treated chamber slides and centrifuged at 1000 × g for 60 s to sediment the condensates. Condensate formation was assessed using fluorescence channels and differential interference contrast (DIC) imaging with a 63× oil-immersion objective (HC PL APO 63×/1.40–0.60 OIL) on a Leica Thunder Imager equipped with a Leica DFC9000 GTC camera. For each well, five random fixed-sized fields of view (2048 × 2048 pixels; pixel size: 0.01 µm^2^) were imaged, and the center 512 × 512 pixels were cropped for display.

### Quantification of Condensates

To measure the total area of coverage, number, and size of condensates, images from fixed-size fields across multiple replicates were processed in CellProfiler (version 4.2; Broad Institute)^102^. Grayscale images were normalized for illumination by applying background correction through the “CorrectIlluminationCalculate” and “CorrectIlluminationApply” modules. Condensates were identified using the “IdentifyPrimaryObjects” module with Otsu’s thresholding to distinguish them from the background, followed by object separation and size filtering to exclude noise and artifacts. The total condensate area was quantified using the “MeasureImageAreaOccupied” module, while the number, size, and circularity of individual condensates were calculated with the “MeasureObjectSizeShape” module. The “MeasureObjectIntensity” module was used to determine each condensate’s total intensity and area, with average intensity per pixel derived by dividing total intensity by area.

### Electrophoretic Mobility Shift Assay (EMSA)

To investigate binding between PAR and FUS, EMSA was performed using Cy3-labeled 16-mer PAR and unlabeled His-MBP-tagged FUS protein. Each 10 µL binding reaction was prepared in a buffer containing 50 mM Tris-HCl (pH 7.5), 100 mM KCl, 10 mM β-mercaptoethanol, 0.1 mg/mL bovine serum albumin (BSA), and 2 mM MgCl₂. The final PAR concentration was 5 nM, while protein concentrations varied depending on the reaction conditions. Samples were incubated at 23 °C for 30 min before being loaded onto a pre-run native 6% TBE polyacrylamide gel (1 mm thick). The gel was prepared 2 h prior to the experiment and pre-run in 0.5× TBE buffer for 30 min.

Electrophoresis was conducted in 0.5× TBE buffer at a constant voltage of 100 V for approximately 60 min. Gels were imaged using a Li-Cor Odyssey M scanner (Serial Number: ODM-0375) with settings optimized for Cy3 fluorescence detection. Data analysis was conducted in Fiji (ImageJ), where background fluorescence from the gel was subtracted, and the intensity of unbound PAR in each lane was quantified and normalized to the no-protein control lane. Equilibrium dissociation constants (K_D_) were calculated by plotting normalized unbound Cy3-PAR intensities against protein concentrations and fitting the data using the Hill equation in GraphPad Prism (version 10.3).

### Fluorescence Anisotropy Assay

To characterize the interactions between PAR and FUS, fluorescence anisotropy assay was used to quantify changes in fluorescence polarization (FP) of PAR upon protein binding. A 4× binding buffer containing 200 mM Tris-HCl (pH 7.5), 400 mM KCl, 40 mM β-mercaptoethanol, 0.4 mg/mL bovine serum albumin (BSA), and 8 mM MgCl_2_ was prepared and diluted to 1× for all dilutions and reaction mixtures. AF488-labeled PAR was diluted to a working concentration of 3 nM in 1× binding buffer. Protein titrations were prepared using 5× stock concentrations, which were serially diluted in 1× binding buffer across 15 steps to achieve final concentrations based on the experimental conditions. For each reaction, 20 µL of PAR master mix and 5 µL of protein dilution were combined in Corning 384-well low-flange black flat-bottom polystyrene microplates (Corning, 3575) to form a 25 µL reaction volume. Negative control wells contained 5 µL of 1× binding buffer instead of protein. Samples were prepared in triplicate to ensure reproducibility. Plates were gently mixed and incubated at 23 °C for 30 min to allow binding equilibration. After incubation, fluorescence polarization was measured using a BMG LABTECH CLARIOstar Plus microplate reader (Serial Number: 430-4336) equipped with SMART software (version V6.20) set to an excitation wavelength of 488 nm and an emission wavelength of 520 nm. Polarization values were analyzed to calculate equilibrium dissociation constants (K_D_) by fitting the data using the Hill equation in GraphPad Prism (version 10.3), providing insights into the binding affinity of the FUS–PAR complexes. Changes in polarization upon PAR saturation by proteins were also used to assess the hydrodynamic properties of PAR during its interaction with FUS.

### Dynamic Light Scattering (DLS) Analysis

Dynamic Light Scattering (DLS) was used to analyze condensates formed by FUS with PAR. Samples were prepared following the protocol described in the ***In Vitro* Reconstitution of Condensates and Microscopy** section, with all buffers filtered through 0.22 µm syringe filters before use. After the addition of TEV protease, samples were incubated in a cuvette for 4 h, and measurements were taken at 1, 5, 15, 30, 60, 90, 120, and 240 min. DLS measurements were performed using a DynaPro Molecular Sizing Instrument (Wyatt Technology) equipped with an 655 nm laser and a temperature-controlled microsampler set to 25 °C. Instrument settings included an acquisition time of 5 s, sensitivity at 90%, and a signal-to-noise threshold of 2.5. An average of 10 acquisitions per time point was used to calculate the mean autocorrelation function. Hydrodynamic radii were derived by resolving the autocorrelation function into Gaussian distributions using Dynamic 6 software (Wyatt Technology). The data were visualized using SigmaPlot.

### Fluorescence Recovery After Photobleaching (FRAP) Analysis

Fluorescence Recovery After Photobleaching (FRAP) experiments were performed using a Leica Thunder Imaging system equipped with an infinity scanner for photobleaching and a Leica DFC9000 GTC camera for fluorescence detection. Condensates containing 10% AF488-labeled FUS and unlabeled PAR were photobleached using 100% 488 nm laser power for 150 iterations. Time-lapse images were acquired starting right after bleaching at 30-s intervals using a 63× oil-immersion objective (HC PL APO 63×/1.40–0.60 OIL) to monitor fluorescence recovery dynamics. Fluorescence intensities were measured within the bleached regions of interest (ROIs), an adjacent unbleached reference region, and a background area using Fiji (ImageJ). Fluorescence intensities in the bleached ROIs were first background-subtracted and then normalized to their pre-bleaching values by calculating the ratio of post-bleaching fluorescence intensities to the initial fluorescence intensity. Recovery parameters, including the mobile fraction and half-time of recovery (T_1/2_), were determined by fitting the recovery curves to a single-exponential equation:

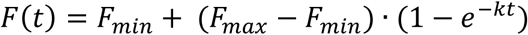

Where:

● 𝐹(𝑡) is the normalized fluorescence intensity at time 𝑡,
● 𝐹_𝑚𝑎𝑥_ is the maximum normalized fluorescence intensity achievable,
● 𝐹_𝑚𝑖𝑛_ is the lowest normalized fluorescence intensity achieved upon photobleaching,
● 𝑘 is the rate constant.

The mobile fraction and half-time of recovery (T_1/2_) were calculated as:

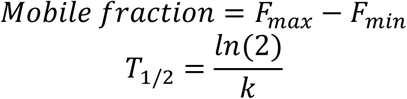

𝑇_1/2_ represents the time 𝑡 when the normalized fluorescence intensity equals 50% of the maximum recovery (i.e., 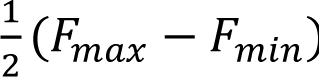)

Control regions within unbleached areas of the same sample were analyzed to account for potential photobleaching and phototoxicity artifacts.

### Limited Proteolysis Mass Spectrometry (LiP-MS)

#### Reaction Preparation

Each condition was prepared in triplicate. HEPES buffer (pH 7.5), NaCl, and FUS protein were combined in a tube with final concentrations of 25 mM, 100 mM, and 0.5 μM, respectively. For conditions containing PAR, 16-mer PAR was added to a final concentration of 0.5 μM. Samples were incubated at room temperature for 2 min before adding 25 units of TEV (final concentration is 4 µM) protease to each sample. Samples were further incubated at room temperature for 2 h.

For the limited proteolysis step, Proteinase K (PK, Thermo Scientific 17916, prepared as a 1 mg/mL stock in 10% glycerol, aliquoted, flash frozen, and stored at −20°C) was diluted to 0.05 mg/mL. A 2 μL aliquot of the diluted PK was placed at the bottom of a new 1.5 mL tube, and 100 μL of the sample was added and mixed rapidly by pipetting up and down seven times. The mixture was incubated for exactly 10 s before quenching the reaction by placing the tube in a 105°C mineral oil bath for 5 min. After cooling and centrifugation, the reaction mixture was transferred to a tube containing solid urea to achieve a final concentration of 8 M urea. Cysteines were reduced with dithiothreitol (DTT) to a final concentration of 10 mM and incubated for 30 min at 37°C with shaking (700 rpm). Subsequently, cysteines were alkylated with iodoacetamide (IAA) at a final concentration of 40 mM for 45 min at room temperature in darkness.

The sample was diluted fourfold with 100 mM ammonium bicarbonate to achieve a final urea concentration of 2 M. Trypsin was added at a 1:50 w/w ratio (trypsin:FUS, 0.2 μg for a 100 μL reaction volume) and incubated overnight at 25°C with shaking at 700 rpm. Digested samples were acidified with trifluoroacetic acid (TFA) to a final concentration of 1%. Samples were desalted using Sep-Pak Vac 1cc (50 mg) C18 cartridges (Waters) and dried using a vacuum centrifuge. Dried peptides were stored at −80°C until ready for mass spectrometry analysis.

#### Mass Spectrometry Acquisition

Peptide samples were resuspended in 0.1% formic acid in LC-grade water to a final concentration of 0.33 μg/μL, and approximately 1 μg of peptides was injected into a Thermo UltiMate 3000 UHPLC system for chromatographic separation. Chromatography solvents included 0.1% formic acid (Solvent A) and 0.1% formic acid in acetonitrile (Solvent B). Peptides were trapped on an Acclaim PepMap 100 C18 column (75 μm × 2 cm, 3 μm, 100 Å) for 10 min at 2% Solvent B, followed by separation on an Acclaim PepMap RSLC C18 column (75 μm × 25 cm, 2 μm, 100 Å) using a 95-min linear gradient from 5% to 25% Solvent B, then a further 25-min gradient from 25% to 40% Solvent B. A Thermo Q Exactive HF-X Orbitrap mass spectrometer was used for analysis, performing a full MS scan at 120,000 resolution followed by 20 data-dependent MS scans at 15,000 resolution.

#### Data Analysis

Label-free quantification (LFQ) was performed using Proteome Discoverer (PD, version 2.4), employing MSFragger and Minora Feature Detector nodes for peptide identification and quantification. Peptide features were searched against a FUS-specific FASTA database, with semi-specific trypsin searches allowing up to two missed cleavages. The Philosopher node validated results with a false discovery rate (FDR) cutoff of 1%. Normalized ion counts were analyzed across replicates, and effect sizes (log2 ratios) and significance (-log10 P values) were calculated using Welch’s t-test.

Missing values were imputed as follows: If a feature was undetected in only one injection across all six injections (three replicates each for two conditions), the missing value was excluded. Features missing in more than one injection across all six injections (e.g., one in each condition or two in one condition) were excluded from analysis. However, if all three replicates in one condition were missing but all three replicates in the other condition were present, the missing values were replaced with the detection limit of the mass analyzer (1000 ion counts).

For peptides detected in multiple charge states, data were consolidated based on median quantification and combined P values using Fisher’s method if consistent in sign; otherwise, the P value was set to 1. Peptides with effect sizes >1 (log2 ratios) and P values >1.3 (-log10 P values) were deemed significant.

#### Cleaved Residue Visualization

Enriched half-tryptic peptides were mapped to Proteinase K (PK) cleavage sites. Fully tryptic peptides were centered to indicate PK accessibility on the opposing side. Data visualization included heat maps, with residue positions plotted on the x-axis. Accessibility changes were represented by spikes proportional to peptide ratios (y-axis) and colored by P value significance (-log10 scale). Plots were generated using Python with Matplotlib, employing normalized ion counts, effect sizes, and P values.

#### Biolayer Interferometry (BLI) Assay

Biolayer interferometry (BLI) experiments were conducted using an Octet RED96 system (Sartorius), using HIS1K biosensors (Sartorius, 18-5120). The HIS1K biosensors were hydrated in PBSA (PBS + 0.1% (w/v) bovine serum albumin (BSA)) for 10 min before loading. Reactions were prepared in black flat-bottom (chimney well) 96-well polypropylene microplates (Greiner Bio-One, 655209). His-MBP-tagged PrLD (400 nM in PBSA) was immobilized onto the biosensor surface during a 120-s loading step, followed by a 60-s baseline equilibration in PBSA. For the association phase, MBP-tagged CTRR was prepared in PBSA at varying concentrations (0.4, 1, 2, 5, and 12 µM) and incubated with the immobilized PrLD for 600 s. For the dissociation phase, the biosensors were transferred to PBSA for 600 s. For PAR competition assays, following the association phase, 16mer-PAR was added to the dissociation buffer at increasing concentrations (0.08, 0.4, 2, and 10 µM) to evaluate its impact on CTRR binding. Regeneration of biosensors was performed using 10 mM glycine (pH 2.7) for 5 s, followed by neutralization in PBSA for 5 s, repeated three times.

Data were processed and analyzed using Octet Data Analysis HT Software version 7.1 (Sartorius), with reference (association and dissociation in buffer alone) subtraction and inter-step correction. Kinetic parameters, specifically the rate of association (𝑘_𝑜𝑛_), the rate of dissociation (𝑘_𝑜𝑓𝑓_), and the equilibrium dissociation constant (𝐾_𝐷_), were derived by fitting the association phase data to an exponential kinetic binding model in GraphPad Prism (version 10.3) using the following equations:

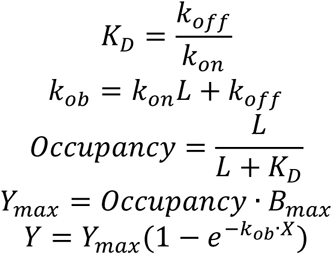

Where:

● 𝐿 is the concentration of analyte (CTRR) in molars (M),
● 𝑋 is time in seconds (s),
● 𝑌 is binding responses in nanometers (nm),
● 𝐵_𝑚𝑎𝑥_ is the maximum binding response achievable in nanometers (nm).

### Single-Molecule Analysis of PAR Conformational Change and PAR–Protein Interactions

#### Single-Molecule Platform Setup

Single-molecule measurements were conducted on a PEG-passivated quartz slide and observed using a home-built prism-type total internal reflection fluorescence microscopy (TIRFM). The TIRFM setup and slide preparation followed protocols from the Ha Lab^103^. To prepare the slides, a 2% (w/v) solution of biotin-PEG (biotin-PEG-5000, Laysan Bio, Inc.) was mixed with m-PEG (m-PEG-5000, Laysan Bio, Inc.) and incubated overnight on amino-silane-treated glass coverslips and quartz slides. This method has been shown to provide a clean background and an optimal density of signals^104,105^.

#### FRET Measurements

To prepare the slide surface for experiments, 1 mg/mL of NeutrAvidin was flowed over the surface and subsequently washed with T50 buffer (10 mM Tris-HCl, pH 7.4, and 50 mM NaCl). Partial duplex PAR probes with Cy3–Cy5 dual labeling, prepared as previously described^51^, were introduced and bound to the biotin-functionalized surface. Specific concentrations of unlabeled FUS or mutants were then added as indicated, and Cy3 and Cy5 signals were tracked and analyzed using IDL and MATLAB scripts established in previous studies^4^. All single-molecule images were captured in an imaging buffer consisting of 20 mM Tris-HCl (pH 7.4), 100 mM KCl, 0.5% (w/v) glucose, 1 mg/mL glucose oxidase, 1.8 U/mL catalase, and 10 mM Trolox.

#### Dwell Time Measurements

For dwell time analysis, slides were prepared as described above. Cy5-labeled partial duplex PAR probes, prepared as previously described^51^, were immobilized on the PEG-passivated surface. Cy3-labeled FUS was then introduced at defined concentrations. Individual Cy3 binding events were localized and tracked over time. The appearance and disappearance of Cy3 fluorescence spikes were quantified to determine dwell times, which were analyzed using the same IDL and MATLAB pipeline^4^. The imaging buffer was identical to that used for FRET measurements.

#### Fluorescence Correlation Spectroscopy (FCS)

Fluorescence correlation spectroscopy (FCS) measurements were performed using a MicroTime 200 system (PicoQuant) equipped with a 532 nm continuous-wave laser for excitation. A 60× water immersion objective (NA 1.2) was used to focus the laser into the sample placed in μ-Slide plastic chambers (ibidi) mounted on a piezoelectric stage. The laser was focused 30 µm above the chamber surface, and the average excitation power at the back aperture of the objective was maintained at ∼5 µW. Emitted fluorescence was filtered using a triple-band filter and a 582/64 nm band-pass filter before detection with single-photon avalanche diodes (SPADs).

The resulting fluorescence intensity traces were autocorrelated to generate autocorrelation curves. Correlation functions were fitted using a standard model for 3D diffusion in a Gaussian-shaped confocal volume:

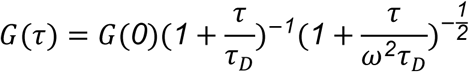

where is 𝜏 the lag time, 𝜏_D_ is the translational diffusion time, ω is the axial-to-lateral dimension ratio of the confocal volume, and 𝐺(*0*) is the autocorrelation amplitude. For confocal volume calibration, we used free Cy3 dye and fitted its autocorrelation curves by fixing its diffusion coefficient 𝐷^𝐶𝑦^^3^ to its known value (241 µm²/s). From this, the structural parameter ω^2^ was extracted using the relationship 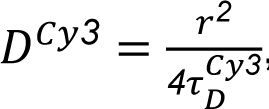, where 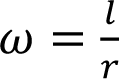.

In subsequent fittings, ω^2^ was held constant.

The hydrodynamic radius 𝑅_𝐻_ was calculated using:

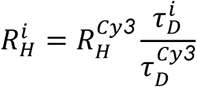

with 𝑅*^cy3^_H_* = 0.81 nm.

Cy3-labeled MBP-FUS variants (20 nM) were used to generate the autocorrelation curves. Given that NHS-based labeling may leave residual free dye, multiple rounds of desalting were performed to remove unbound dye. All dilutions were made with nuclease-free water, and each measurement was conducted for ∼180 s. All measurements were repeated at least three times.

#### Cellular Laser Microirradiation and Quantification

U2OS cells were cotransfected with PARP1-chromobody (Chromotek, XCR) and wild-type or mutant GFP-FUS constructs using the JetPrime reagent (Polyplus, 101000046), following the manufacturer’s instructions. Media were replaced 4–6 h after transfection. The next day, cells were split and seeded into 35-mm FluoroDishes with a 0.17-mm coverslip bottom (World Precision Instruments, FD35-100). On the third day, cells were treated overnight with either 1 µM PARG inhibitor PDD 00017273 (PARGi) or DMSO as control. To prepare cells for microirradiation, they were pulsed with 200 ng/mL Hoechst 33342 (Thermo Scientific, Cat# 62249) for 30 min, rinsed once with PBS, and then phenol-free FluoroBright DMEM (Gibco, A18967-01) was added for imaging.

Laser microirradiation was performed using a 355-nm OPSL laser (Coherent, Genesis). Time-lapse images, including pre- and post-irradiation frames, were captured using a Zeiss LSM880 laser-scanning confocal microscope with a 40× water-immersion objective (N.A. 1.2, C-Apochromat).

For quantitative evaluation of recruitment kinetics, fluorescence intensities of the irradiated regions (ROI) were measured using Fiji (ImageJ) and normalized to pre-irradiation values of the same regions later being quantified as DNA damage lines. Fluorescence intensities from non-irradiated regions in the same cells were used as background controls. Corrected total cell fluorescence (CTCF) was calculated using the formula (https://theolb.readthedocs.io):

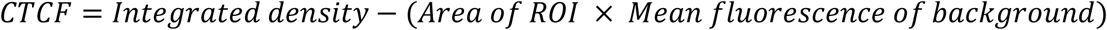

### Molecular Dynamics (MD) Simulations

#### Modeling of FUS Protein Using Coarse-Grained (CG) Models

We model the FUS protein using two well-established coarse-grained (CG) models: the native-structure-based Gō-like mode^l^^45,46^ for the structured domains and the hydrophobic scale (HPS) mode^l^^47–49^ for the disordered regions.

#### Gō-Like Model for Structured Domains of FUS

To model the structured domains of FUS, specifically the RNA Recognition Motif (RRM) and the Zinc Finger (ZnF), we employ the Gō-like model developed by Karanicolas and Brooks III^46^. This model incorporates the following components:

● Bonded interactions: Harmonic potentials on bonds and angles, along with a sequence-dependent statistical potential on dihedral angles between adjacent residues.
● Favorable nonbonded interactions: Applied between residue pairs that are in contact in the native structure.
● Repulsive nonbonded interactions: Applied between all other residue pairs not in native contact.

Native contacts are defined based on backbone hydrogen bonds and side-chain interactions observed in the native structure. Contacts between residues separated by fewer than two positions in the primary sequence are excluded.

#### Nonbonded Interactions

The nonbonded interaction energy for residues in native contact is:

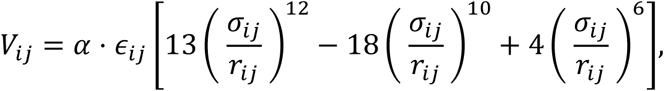

where 𝑟_𝑖𝑗_ is the distance between the two residues, 𝜎_𝑖𝑗_ is the distance at which the interaction energy is minimized, and -𝛼 ⋅ 𝜖_𝑖𝑗_ is the interaction strength at 𝑟_𝑖𝑗_ = 𝜎_𝑖𝑗_. The value of 𝜎_𝑖𝑗_ is set to the distance between the two residues’ 〈 carbon atoms in the native structure. The value of 𝜖_𝑖𝑗_ is the contact energy reported by Miyazawa and Jernigan^106^ based on the types of the two residues and 〈 is a scaling factor applied to all residue pairs in native contact. The value of 〈 is determined by setting the folding temperature of the structured domain. Although the original Gō-like model sets the folding temperature to 350 K by default, here we set the folding temperature to 500 K. Once the value of 〈 is determined, the Gō-like model defines the native contact energy per residue as 𝜖_𝑟𝑒𝑠_ = 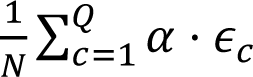, where Q is the total number of native contacts and 𝜖 is the value of the 𝜖 parameter for contact c. The Gō-like model uses 𝜖_𝑟𝑒𝑠_ to define the overall energy scale, based on which other interactions are defined.

Nonbonded interactions for residues that are not in native contact and are separated by more than three residues, are purely repulsive:

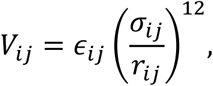

where 𝜎_𝑖𝑗_ is equal to the average of the repulsive radii of residues involved and 𝜖_𝑖𝑗_ = 1.5 × 10^−3^𝜖_𝑟𝑒𝑠_. The repulsive radius of a residue is defined as the distance from its closet residue that is not in native contact.

#### Bonded Interactions

The bonded interactions in the Gō-like model include energy terms for bonds, angles, and dihedrals, that are defined based on the protein’s primary sequence. Both the bond and angle energy terms are harmonic potentials with their equilibrium values set to the values in the native structure. The force constants for the bond and angle potentials are set to 200𝜖_𝑟𝑒𝑠_ and 40𝜖_𝑟𝑒𝑠_, respectively. The dihedral energy term is a cosine series with up to 4 terms, with the parameters set to reproduce the distribution of dihedral angles among the residues in the Protein Data Bank. In other words, the dihedral energy term is sequence-specific and does not depend on the native structure.

For both the RRM and ZnF domains, we generated the Gō-like model using the MMTSB^107^ Gō model builder server (https://mmtsb.org/webservices/gomodel.html). The native structures of the RRM and ZnF domains were obtained using AlphaFold2^108^.

#### Hydrophobic Scale (HPS) Model for Disordered Regions of FUS

The disordered regions are modeled using the hydrophobic scale (HPS) model developed by Dignon^48^ which includes:

● **Bonded Interactions**: Harmonic potentials for bonds between consecutive residues, with a force constant of 8033 kJ/(mol·nm²) and an equilibrium bond length of 0.38 nm.
● **Electrostatic Interactions**: The Debye-Hückel theory is used to model the electrostatic interactions as

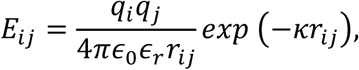

where 𝑞_𝑖_ and 𝑞_𝑗_ are net charges of residue *i* and *j,* 𝜖_0_ is the dielectric constant of vacuum, 𝜖_𝑟_ = 80 represents the relative dielectric constant of water, and κ^-1^ is the Debye length determined by the ionic effects on electrostatic screening, with 𝜅^−1^ = 0.304/√𝐼 nm, where 𝐼 is the solution’s ionic strength in molarity.

- ● **Short-Range Van der Waals Interactions**: We use the follow to model the short-range Van der Walls interactions:

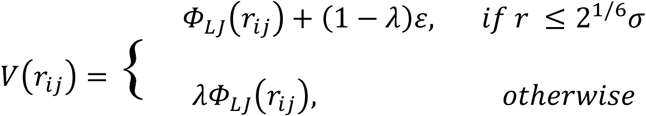

where 𝛷_𝐿𝐽_(𝑟) = 4𝜀[(𝜎/𝑟)^12^ − (𝜎/𝑟)^6^] and ε = 0.8368 kJ/mol. σ and λ are the mean of amino acid-specific parameters quantifying their size and hydrophobicity, respectively. We use the values from the M1 model developed by Tesei et al.^49^ for σ and λ of protein residues.

#### Customized HPS Modeling for PAR Based on smFRET Measurements

To model PAR, we developed a customized HPS model based on previous single-molecule Förster resonance energy transfer (smFRET) measurements of the end-to-end distance of PARs with various lengths^51^. Each ADP-ribose unit is represented by a single bead, with a harmonic bond potential between consecutive ADP-ribose units. The equilibrium bond length is 1.16 nm, matching the monomeric ADP-ribose length, and the force constant is 8033 kJ/(mol•nm²). The net charge of each ADP-ribose unit is -2 due to the diphosphate group. We optimized the parameters σ and λ of short-range van der Waals interactions between ADP-ribose units by fitting them to the experimental FRET data of PARs with different lengths. Using the conformations from the simulations, we calculated the average FRET efficiency based on the FRET theory:

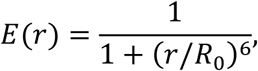

where R_0_ = 5.9 nm is the Förster distance, r is the end-to-end distance of a PAR conformation, and E(r) is the FRET efficiency at distance r. By comparing the computed average FRET efficiency with experimental data, we determined the optimal values of σ and λ (1.5 nm and 0.8, respectively), resulting in simulated FRET efficiencies that closely matched the experimental data.

#### Slab Simulation of FUS–PAR Condensation

We conducted slab simulations to investigate FUS condensation in the presence and absence of 16-mer PAR. For the FUS-only simulation, the system contained 100 FUS proteins. In the FUS–PAR simulation, we matched experimental concentration ratios by including 100 FUS proteins and 10 PAR molecules.

The initial configurations were prepared as follows:

● **System Setup**: FUS proteins (and PAR molecules, if applicable) were positioned in an elongated simulation box with dimensions of 25 nm × 25 nm × 3500 nm along the x, y, and z axes. The large z-axis dimension ensured adequate spacing, preventing clashes.
● **Equilibration**: The system was equilibrated with a 100 ns NVT simulation at 250 K, using a Langevin integrator and a time step of 10 fs.
● **Compression**: Following equilibration, a 300 ns NPT simulation was conducted at 250 K and 1 atm, using an anisotropic Monte Carlo barostat that adjusted only the z-axis dimension. This compression along the z-axis produced a condensed phase, which served as the initial configuration for slab simulations.
● **Production Slab Simulations**: The production runs were conducted in an NVT ensemble in a periodic box with dimensions of 25 nm × 25 nm × 200 nm (x, y, z). Each slab simulation was run for 5 μs with a time step of 10 fs. Simulations for the FUS-only system were performed at temperatures between 330 K and 420 K, while the FUS–PAR system extended from 330 K to 440 K.

#### Computation of the Critical Temperature (*T_c_*) for Condensation from Slab Simulations

To determine the critical temperature (*T*_c_) for condensation, we analyze results from slab simulations using the same method proposed by Dignon et al^48^. At a given temperature (*T*), we assume that the density difference between the dense (ρ_H_) and dilute (ρ_L_) phases follows the relationship:

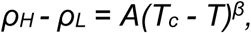

where the critical exponent β is set to 0.325. We fit the system-specific parameters A and *T*_c_ using ρ_H_ and ρ_L_ values obtained from slab simulations across various temperatures.

#### Steps to Calculate ρ_H_, ρ_L_, and the Density Profile along the z-axis

- **Cluster Analysis**: For each simulation frame, we perform hierarchical clustering of the FUS proteins, using single-linkage clustering based on the pairwise distances of their centers of mass. Clusters are defined by setting a dendrogram cutoff at 5 nm.
- **Centering**: Each frame is centered by shifting the center of mass of the largest cluster to z = 0 to standardize the position of the dense phase along the z-axis.
- **Density Profile Computation**: To compute the density profile along the z-axis, the simulation box is divided into 60 equal-sized bins along this axis. We calculate the density in each bin.
- Determining ρ_H_ and ρ_L_:
- ρ_L_ is calculated as the average density in the regions where ∣z∣ > 50 nm.
- ρ_H_ is calculated as the average density in the regions where ∣z∣ < 5 nm.
- These density values are averaged over all simulation frames to obtain consistent ρ_H_ and ρ_L_ values.

#### Contact Frequency Analysis

The second half of the simulation trajectory (2.5–5.0 µs; 25,000 frames) was analyzed to generate contact frequency maps between PAR and FUS residues, as well as among FUS residues. Contacts were defined using a distance cutoff of 1.2 nm. A FUS residue was considered to be in contact with PAR if it was within 1.2 nm of any PAR bead. A FUS protein was considered to be in contact with PAR if any of its residues met this criterion. Similarly, two FUS residues were considered in contact if the distance between them was below the cutoff. Contact frequencies were computed by averaging over simulation frames and all FUS molecules.

For the PAR–FUS contact analysis, we accounted for the fact that there are more FUS molecules than PAR chains in the simulation box, meaning not all FUS proteins interact with PAR at all times. To assess the relative contact frequencies of different FUS regions with PAR, we excluded frames in which a FUS protein made no contact with PAR. The average contact frequency for each FUS protein was calculated using only the retained frames. These values were then averaged across all FUS proteins to yield the final relative contact frequency profile of FUS residues with PAR.

To investigate how PAR interactions influence FUS conformations, we computed intra-and inter-residue contact frequency maps for FUS both in the presence and absence of PAR. For the FUS-only system, contact frequencies between FUS residues were calculated by averaging over all simulation frames and FUS proteins. In the FUS–PAR simulations, contact maps were generated using only the frames in which a given FUS protein had at least one residue contacting PAR. The resulting contact frequencies were then averaged over all FUS proteins. These contact maps were compared to those from the FUS-only simulations by calculating the ratio of contact frequencies in the presence versus absence of PAR.

#### Computation of the Thermodynamics of Binding with Umbrella Sampling

We used umbrella sampling^109^ to calculate the binding free energies both between PAR and the FUS domains, as well as among FUS domains. In each umbrella sampling window, an external restraining potential is added to control the distance between the two molecules. It is defined as:

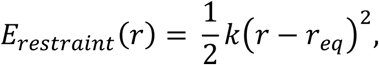

where *k =* 50 kJ/(mol·nm²) is the force constant, *r* represents the distance between the centers of mass of the two molecules, and *r_eq_* is the equilibrium distance. We conducted simulations across 61 windows, with *r_eq_* linearly varying from 0 to 20 nm.

#### Simulation Protocol

● **Initial Configurations**: Each window was energy-minimized before starting the simulations.
● **Equilibration**: We performed a 100 ns equilibration in the NVT ensemble at 293.15 K.
● **Production**: Following equilibration, a 30 µs production run was conducted in the NVT ensemble at the same temperature, with a time step of 10 fs for both equilibration and production phases.

Using data from all windows, we constructed the potential of mean force (PMF) along *r* using the multistate Bennett acceptance ratio (MBAR) method^110^, as implemented in the FastMBAR package^111^. From the PMF, we calculated the binding free energy by classifying configurations into bound and unbound states, using a cutoff distance of 10 nm. To estimate uncertainty in the binding free energy, the trajectory from the last 25 µs was divided into five segments, each used independently to calculate the binding free energy, providing a basis for estimating the uncertainty.

## SUPPLEMENTAL INFORMATION

Supplemental Information includes five figures and a movie.

## ACKNOWLEDGMENTS

We thank Drs. Phil Sharp, Geraldine Seydoux, Danfeng Cai, Hyun O. Lee, Tatjana Trcek, Lingyao Wang and Hue Sun Chan; Christopher Chin Sang and Genzhe Lu; Dr. Shang-Jung Cheng and all the members of the Leung lab; and ReVision: A Scientific Editing Network at Johns Hopkins University for their valuable feedback on the manuscript. We are also grateful to Drs. Priya R. Banerjee and Gable M. Wadsworth for providing the synthetic polypeptide [RGRGG]₅ and its variants. We thank Dr. Morgan Dasovich, Frances Middleton-Davis, Sarah M. Voss, and Shuaichen Liu for assistance with preliminary work. Schematic models were created using BioRender.com. Simulation work was performed using the Tufts University High Performance Compute Cluster (https://it.tufts.edu/high-performance-computing).

This work was supported by the National Institutes of Health (NIH) under grants T32-GM149382 (H.E.T.), T32-CA009110 (M.B.), R35-GM153387 (M.T.B.), R01-EY031097 (J.B.S.), RF1-NS113636 (S.M.), and R01-AG071326 (A.K.L.L. and S.M.); and by the National Science Foundation (NSF) through CAREER Award 2143160 (J.B.S.). Development of the smFRET probe and PAR labeling was supported by NIH grant R01-GM104135 (A.K.L.L.). H.L. is supported by the Cross-Disciplinary Graduate Program in Biomedical Sciences (XDBio) Fellowship from the Johns Hopkins School of Medicine.

## AUTHOR CONTRIBUTIONS

Conceptualization, H.L., M.B., S.D.F., X.D., and A.K.L.L.; Methodology, H.L., L.S., M.P., N.D., H.E.T., Y.G., M.B., M.T.B., S.M., S.D.F., X.D., and A.K.L.L.; Software, K.Y., and X.D.; Validation, H.L., Y.C., L.S., M.P., N.D., H.E.T., Y.G., K.Y., and X.Y.; Formal Analysis, H.L., Y.C., L.S., M.P., N.D., H.E.T., Y.G., K.Y., and X.D.; Investigation, H.L., Y.C., L.S., M.P., N.D., H.E.T., Y.G., K.Y., X.Y., C.S.F., and X.D.; Resources, J.B.S., M.T.B., S.M., S.D.F., X.D., and A.K.L.L.; Data Curation, H.L., Y.C., L.S., M.P., N.D., H.E.T., Y.G., K.Y., and X.Y.; Writing – Original Draft, H.L., X.D., and A.K.L.L.; Writing – Review & Editing, H.L., M.P., H.E.T., K.Y., M.B., J.B.S., M.T.B., S.M., S.D.F., X.D., and A.K.L.L.; Visualization, H.L., Y.C., N.D., K.Y., and X.D.; Supervision, H.L., M.T.B., S.M., S.D.F., X.D., and A.K.L.L.; Project Administration, H.L., M.T.B., S.M., S.D.F., X.D., and A.K.L.L.; Funding Acquisition, H.L., M.B., J.B.S., M.T.B., S.M., S.D.F., X.D., and A.K.L.L.

## DECLARATION OF INTERESTS

The authors declare no competing interests.

---

**Figure S1, related to Figure 1.**
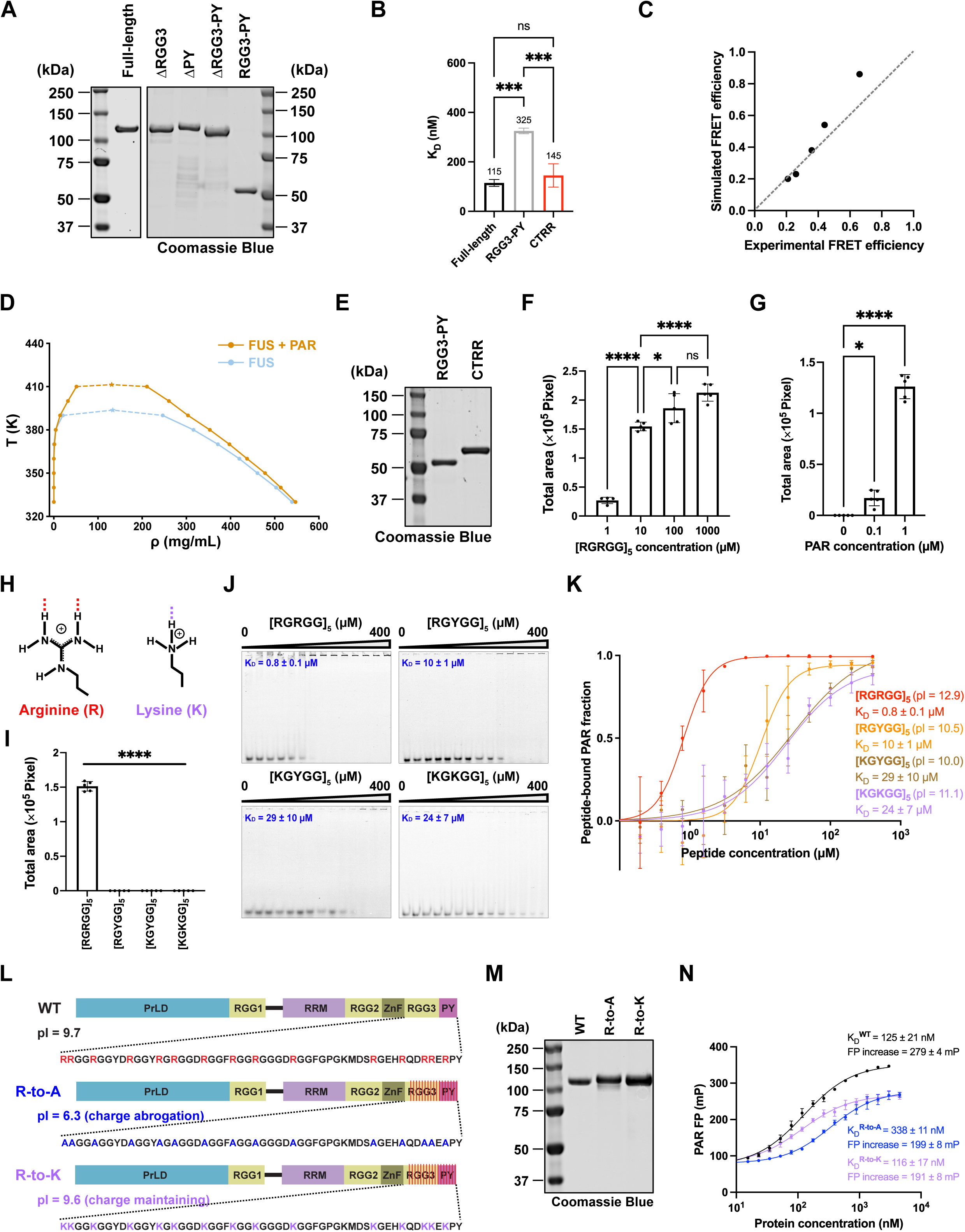
**(A)** Representative SDS-PAGE (4% stacking gel and 10% resolving gel) with Coomassie Brilliant Blue staining showing the purified His-MBP-tagged full-length FUS and its mutants, as presented in Figure 1. The His-MBP solubility tag is specifically cleaved by tobacco etch virus (TEV) protease to initiate the condensation reaction. **(B)** Comparisons of K_D_ values in Figure 1E. n = 3; error bars, SD; ordinary one-way ANOVA with Tukey’s multiple comparisons test, not significant (ns) and p < 0.001 (***). **(C)** Correlation plot comparing FRET efficiency between simulations and experiments for different PAR lengths (see **Methods**). **(D)** Phase diagram of FUS condensation with and without PAR from slab simulations. The densities of the low-density phase (ρ_L_, points on the left) and high-density phase (ρ_H_, points on the right) are plotted against temperature. The critical temperature, where the two phases become indistinguishable (indicated by the top of each curve), is extrapolated from data points at various temperatures (see **Methods**). The critical temperature is approximately 394 K for FUS alone (blue curve) and approximately 411 K for the FUS + PAR system (amber curve). **(E)** Representative SDS-PAGE (4% stacking gel and 12% resolving gel) with Coomassie Brilliant Blue staining showing the purified His-MBP-tagged RGG3-PY fragment and the CTRR fragment, as presented in Figure 1. **(F)** Bar graph quantification of the total area covered by condensates formed in Figure 1H. n = 5; error bars represent SD; ordinary one-way ANOVA with Tukey’s multiple comparisons test, with significance levels: not significant (ns), p < 0.05 (*), and p < 0.0001 (****). **(G)** Bar graph quantification of the total area covered by condensates formed in Figure 1I. n = 5; error bars represent SD; ordinary one-way ANOVA with Tukey’s multiple comparisons test, with significance levels: p < 0.05 (*), and p < 0.0001 (****). **(H)** Schematic of arginine and lysine side chains showing charge and potential hydrogen bond formation. **(I)** Bar graph quantification of the total area covered by condensates formed in Figure 1J. n = 5; error bars represent SD; ordinary one-way ANOVA with Tukey’s multiple comparisons test, p < 0.0001 (****). **(J)** Representative electrophoretic mobility shift assay (EMSA) gels showing binding of titrated [RGRGG]_5_ peptide and its arginine-substituted derivatives with 5 nM Cy3-PAR. n = 3. K_D_ values were determined by quantifying unbound Cy3-PAR (the bands at the bottom of the gels) intensities across peptide titrations and fitting the data using a one-site specific binding model. See also (K). **(K)** EMSA binding curves showing the fraction of bound PAR as a function of peptide concentration for [RGRGG]_5_ peptide and its arginine-substituted derivatives, as in (J). Isoelectric points (pIs) are indicated for each peptide. K_D_ values were determined by fitting the data using the Hill equation. n = 3; error bars, SD. **(L)** Domain structures of FUS with the RGG3-PY sequence highlighted, showing arginine substitutions with alanines (R-to-A) or lysines (R-to-K) to eliminate or maintain charge. The theoretical isoelectric points (pIs) are indicated for each protein. **(M)** Representative SDS-PAGE (4% stacking gel and 10% resolving gel) with Coomassie Brilliant Blue staining showing the purified His-MBP-tagged full-length FUS and its R-to-A and R-to-K mutants. **(N)** Fluorescence anisotropy assays of wild-type FUS and its arginine mutants with 3 nM AF488-labeled PAR. Fluorescence polarization (FP) values of AF488-PAR are plotted and fitted by the Hill equation. n = 3; error bars, SD.

**Figure S2, related to Figure 2.**
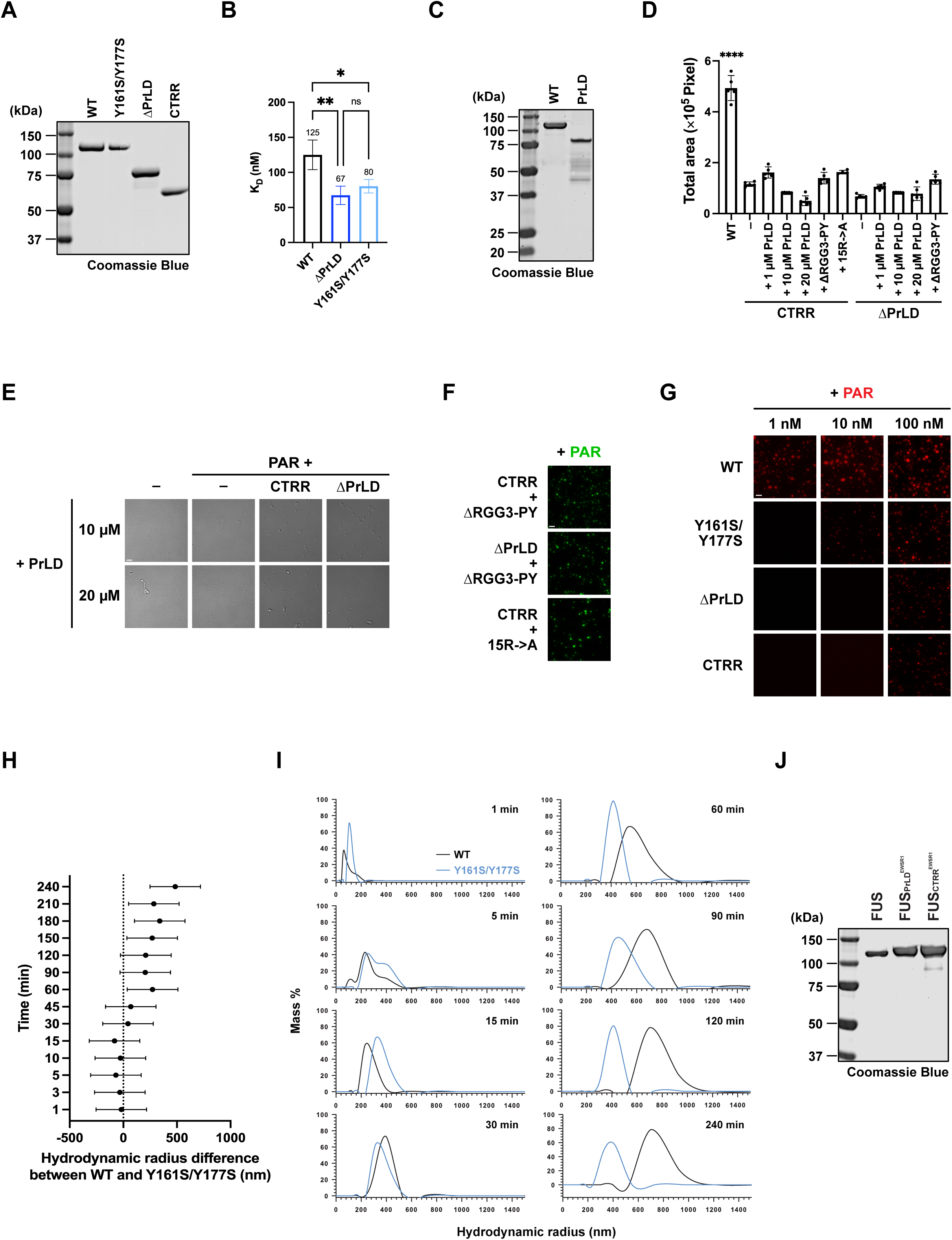
**(A)** Representative SDS-PAGE (4% stacking gel and 10% resolving gel) with Coomassie Brilliant Blue staining showing the purified His-MBP-tagged full-length FUS and its mutants, as presented in Figure 2A. **(B)** Bar graph comparing K_D_ values from Figure 2F. n = 3; error bars, SD; ordinary one-way ANOVA with Tukey’s multiple comparisons test, with significance levels: not significant (ns), p < 0.01 (**), and p < 0.001 (***). **(C)** Representative SDS-PAGE (4% stacking gel and 12% resolving gel) with Coomassie Brilliant Blue staining showing the purified His-MBP-tagged full-length FUS and its PrLD. **(D)** Bar graph quantification of the total area covered by condensates formed in Figures 3G, S3E and S3F. n = 5; error bars represent SD; ordinary one-way ANOVA with Tukey’s multiple comparisons test, p < 0.0001 (****) between wild-type FUS and all the other conditions. **(E)** Representative DIC microscopic images of condensates formed by adding 10 or 20 µM PrLD with the indicated additives. When included, CTRR and ΔPrLD are at 1 µM, and PAR is at 100 nM. n = 5. Scale bar, 5 µm. See also (D). **(F)** Representative fluorescence microscopic images showing condensation results for 1 µM of each protein adding in *trans* with 100 nM PAR. Fluorescence is from Cy3-labeled PAR. n = 5. Scale bar, 5 µm. See also (D). **(G)** Fluorescence microscopy images showing Cy3-PAR signal corresponding to Figure 2H. n = 5. Scale bar, 5 µm. **(H)** Forest plot showing the hydrodynamic radius differences between condensates formed by wild-type FUS and the Y161S/Y177S mutant with PAR at different time points, corresponding to the two-way ANOVA analysis in Figure 2J. n = 3; error bars represent Šidák-adjusted 95% confidence intervals (intervals including 0 indicate no significant difference). **(I)** Representative dynamic light scattering (DLS) mass distribution plots by hydrodynamic radius at the indicated time points, corresponding to Figure 2K. Wild-type FUS and the Y161S/Y177S mutant are compared at each time point. **(J)** Representative SDS-PAGE (4% stacking gel and 10% resolving gel) with Coomassie Brilliant Blue staining showing the purified His-MBP-tagged FUS and its chimeric variants with EWSR1, as presented in Figure 2P.

**Figure S3, related to Figure 3.**
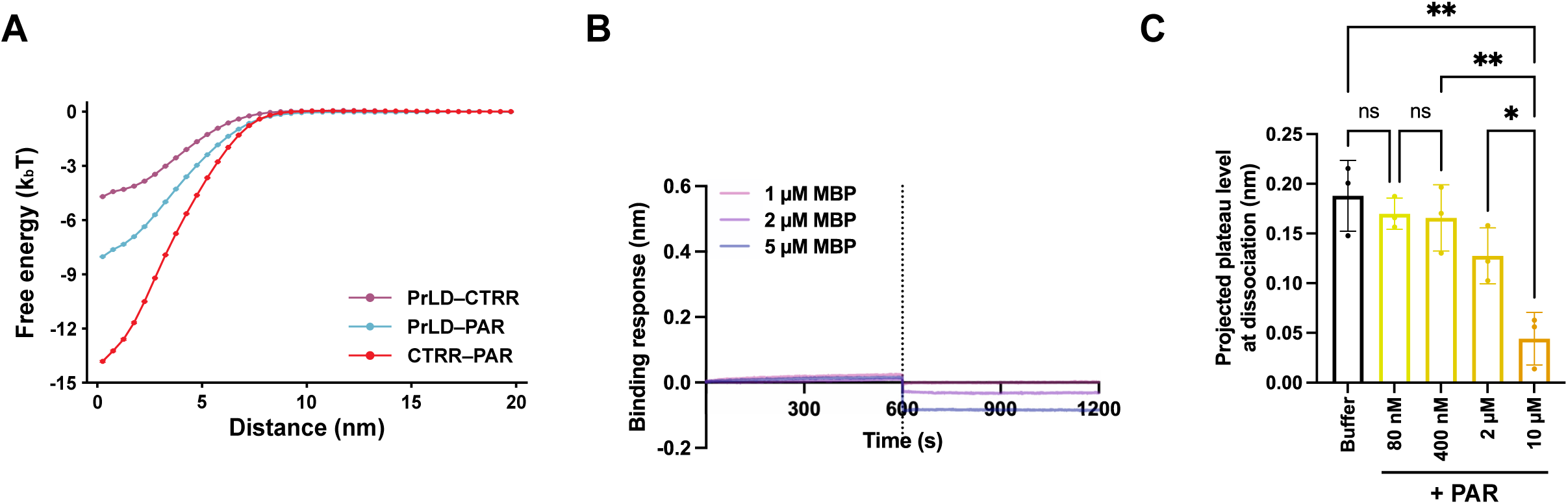
**(A)** Free energy profiles for interactions between PrLD, CTRR, and PAR in umbrella sampling simulations, corresponding to the calculated binding free energies shown in the bar chart in Figure 3E (see **Methods**). n = 5; error bars, SD. **(B)** Representative BLI sensorgram showing no positive binding responses of PrLD and MBP. n = 3. **(C)** Comparison of the projected plateau levels at dissociation by the introduction of buffer and titration of PAR, corresponding to Figure 3H. n = 3; error bars, SD; ordinary one-way ANOVA with Tukey’s multiple comparisons test, with significance levels comparing the projected plateau level at dissociation: not significant (ns), p < 0.05 (*) and p < 0.01 (**).

**Figure S4, related to Figure 4.**
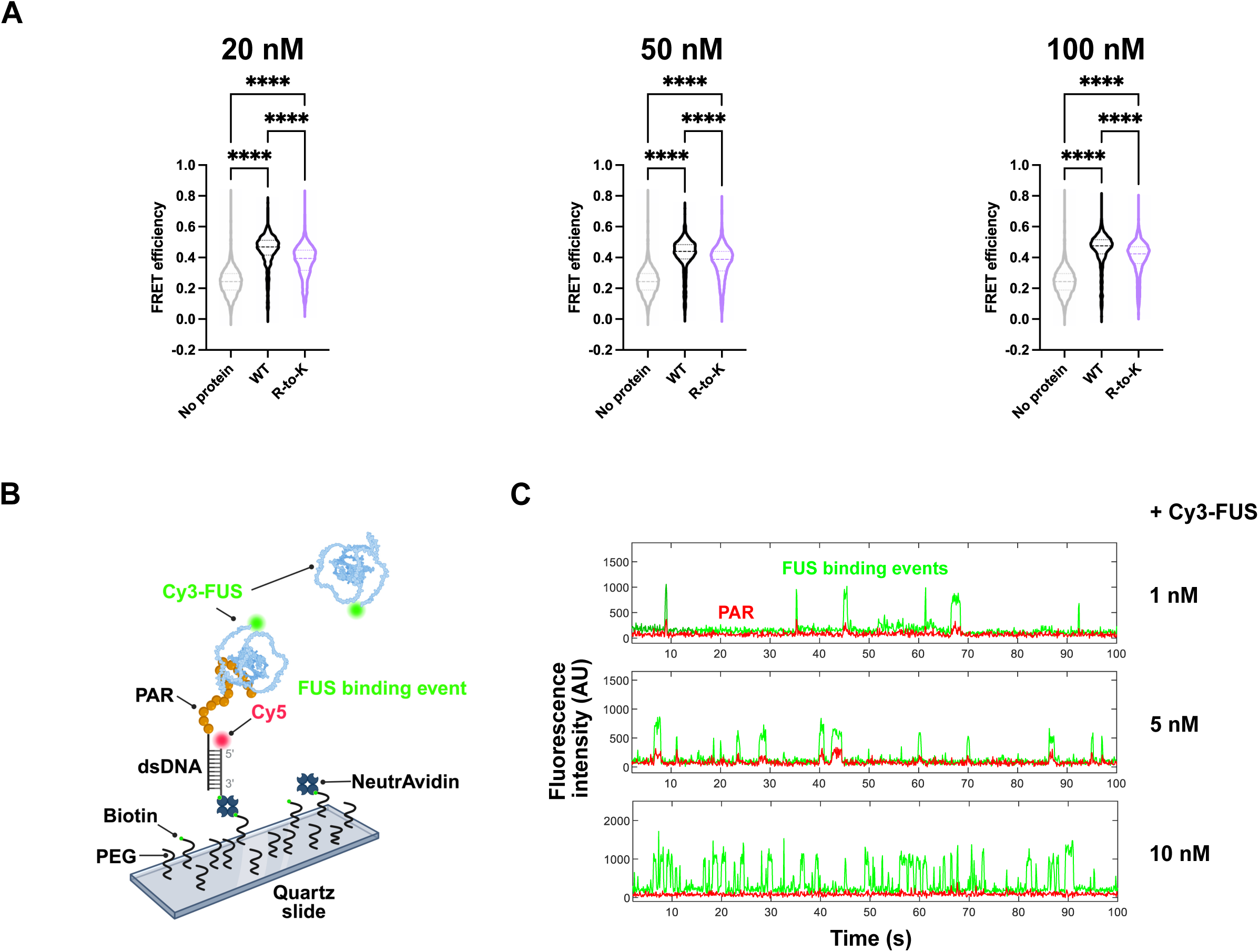
**(A)** Violin plots of PAR FRET efficiencies in response to protein concentrations of 20 nM, 50 nM, and 100 nM. n > 800; dashed lines represent the 75th percentile, the median (50th percentile), and the 25th percentile; Kruskal-Wallis test, p < 0.0001 (****). **(B)** Schematic of single-molecule total internal reflection fluorescence (TIRF) microscopy measurement of Cy3-FUS binding to immobilized PAR, with Cy5-labeled double-stranded DNA (dsDNA) serving as a scaffold. FUS binding to PAR can be detected by the increase in Cy3 signal. **(C)** Representative single-molecule fluorescence traces of Cy3 (green, FUS) and Cy5 (red, immobilized PAR) at increasing Cy3-FUS concentrations. Cy3 signal bursts correspond to transient FUS binding events.

**Figure S5, related to Figure 5.**
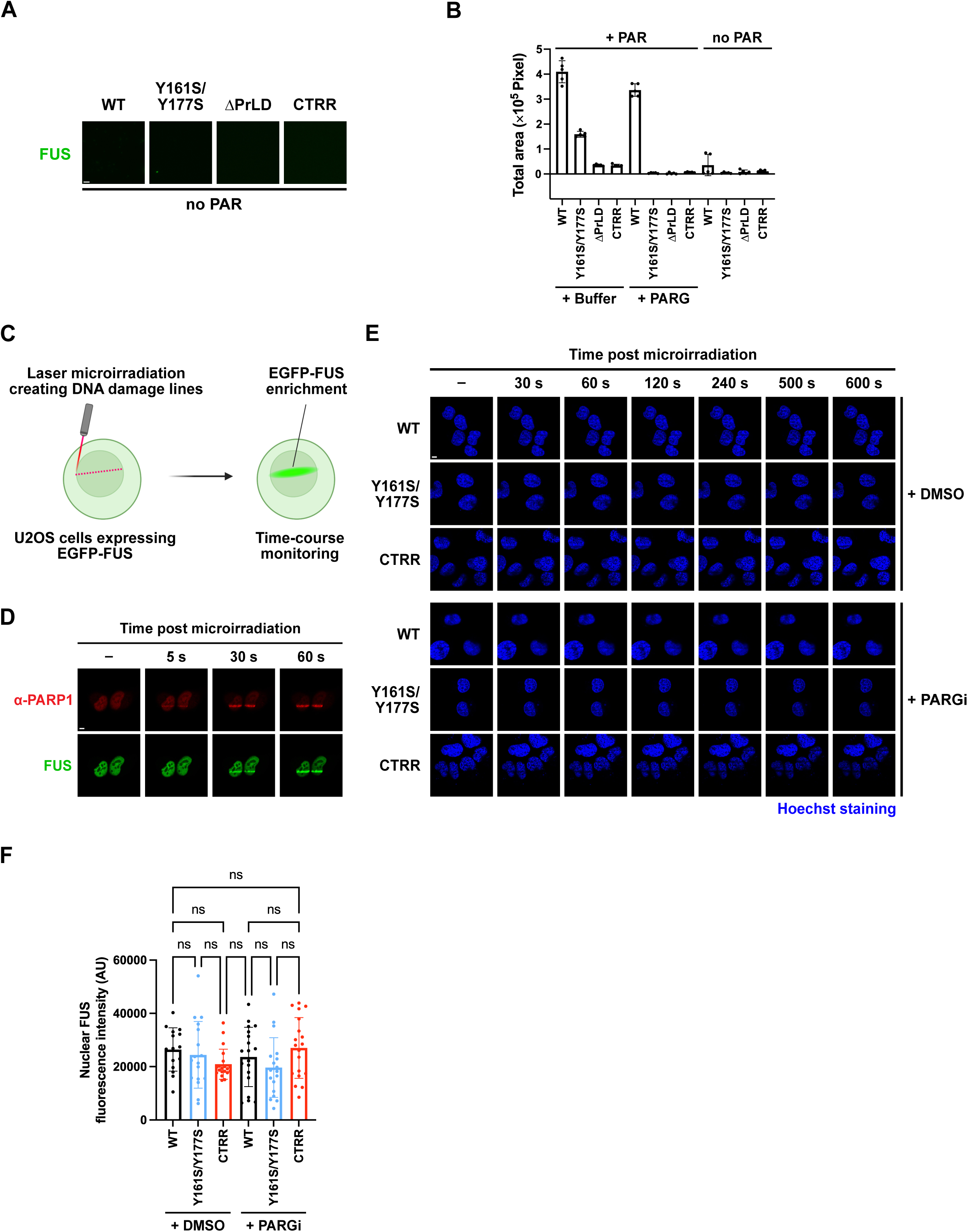
**(A)** Representative microscopic images of condensates formed by 1 µM of wild-type FUS or corresponding mutants (green) without PAR after 2 h incubation. n = 5. Scale bar, 5 µm. See also (B). **(B)** Quantification of the total area covered by condensates formed in Figure 5J and S5A. n = 5; error bars, SD. **(C)** Schematic of the laser strip assay conducted in Figures 5K and 5M. Briefly, a 355 nm ultraviolet-A pulsed laser was used to microirradiate U2OS nuclei transiently transfected with EGFP-FUS, with time-course monitoring of FUS enrichment. **(D)** Representative time-course microscopic images of laser strip assays showing the early recruitment of endogenous PARP1 (red; detected using an RFP-tagged anti-PARP1 chromobody) and the subsequent enrichment of FUS within 1 min post-microirradiation. n = 5. Scale bar, 5 µm. **(E)** Hoechst staining indicating cell nuclei in the laser strip assays, corresponding to Figures 5K and 5M. Scale bar, 5 µm. **(F)** Bar graph showing the nuclear fluorescence intensity of FUS before microirradiation in wild-type FUS, the Y161S/Y177S mutant, and CTRR-expressing cells treated with either DMSO or PARG inhibitor. Each dot represents an individual cell. n ≥ 15; error bars, SD; ordinary one-way ANOVA with Tukey’s multiple comparisons test, showing no significant differences (ns) between any conditions.

## Notes

### Competing Interest Statement

The authors have declared no competing interest.

