## Supplementary figures and images for "Transient Poly(ADP-Ribose) Triggers FUS Condensation Hysteresis via a Prion-Like Mechanism"

### Movie S1

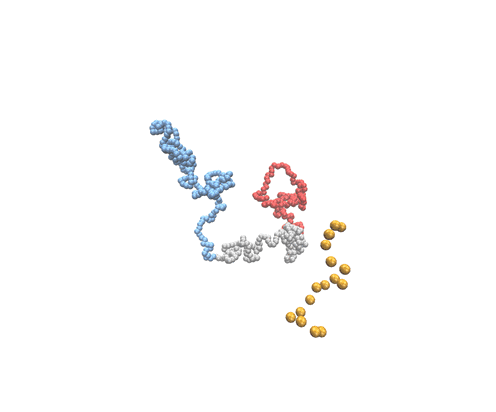
